# Allelic variants of the NLR protein Rpi-chc1 differentially recognise members of the *Phytophthora infestans* PexRD12/31 effector superfamily through the leucine-rich repeat domain

**DOI:** 10.1101/2020.12.29.424692

**Authors:** Daniel Monino-Lopez, Maarten Nijenhuis, Linda Kodde, Sophien Kamoun, Hamed Salehian, Kyrylo Schentsnyi, Remco Stam, Anoma Lokossou, Ahmed Abd-El-Haliem, Richard GF Visser, Jack H Vossen

## Abstract

- *Phytophthora infestans* is a pathogenic oomycete that causes the infamous potato late blight disease. Resistance (*R*) genes from diverse *Solanum* species encode intracellular receptors that recognize *P. infestans* RXLR effector proteins and provide effective defence responses. To deploy these *R* genes in a durable fashion in agriculture, we need to understand the mechanism of effector recognition and the way the pathogen evades recognition.
- We cloned sixteen allelic variants of the *Rpi-chc1* gene from *Solanum chacoense* and other *Solanum* species, and identified the cognate *P. infestans* RXLR effectors. These tools were used to study receptor-ligand interactions and co-evolution.
- Functional and non-functional alleles of *Rpi-chc1* encode Coiled-Coil-Nucleotide Binding-Leucine-Rich-Repeat (CNL) proteins. *Rpi-chc1.1* recognised multiple PexRD12 (AVRchc1.1) proteins while *Rpi-chc1.2* recognised multiple PexRD31 (AVRchc1.2) proteins, both from the PexRD12/31 superfamily. Domain swaps between Rpi-chc1.1 and Rpi-chc1.2 revealed that overlapping subdomains in the LRR were responsible for the difference in effector recognition.
- This study showed that *Rpi-chc1.1* and *Rpi-chc1.2*, evolved to recognize distinct members of the same PexRD12/31 effector family via the LRR domain. The biased distribution of polymorphisms suggests that exchange of LRRs during host-pathogen co-evolution can lead to novel recognition specificities. These insights will help future strategies to breed for durable resistant varieties.

## Introduction

Potato (*Solanum tuberosum*) is the fourth largest food crop in the world after maize, rice and wheat, with more than 368 million tonnes produced in 2018 (FAO, 2020). Potato late blight, caused by the oomycete *Phytophthora infestans* (*P. infestans*), is one of the most infamous potato diseases. During the mid-1840s, this pathogen caused the Great Irish Famine from which around one million people died (Callaway, 2013). Nowadays, losses from late blight are estimated to still reach 16% of the world production and the main disease management is based on biocide applications. Including yield losses and crop protection measures, late blight causes a global economic loss of € 5.2 billion per year (Haverkort *et al.*, 2016).

*P. infestans* is an oomycete with sexual and asexual life cycles, which exhibits a hemibiotrophic lifestyle on potato. Together with its large and fast evolving genome, it leads to the regular emergence of new aggressive and virulent strains. The infection starts when a spore lands on the plant surface, germinates and forms a penetration structure called appressorium. Alternatively, spores can also enter through natural openings such as stomata. After passing the epidermis, hyphae spread intercellularly projecting haustorium structures into the mesophyll cells. These haustoria are specialised infection structures that create an intimate association with the host cell facilitating nutrient uptake, and both apoplastic and cytoplasmic effector secretion (Fry, 2008). Effectors are pathogen molecules that interact with different host targets to suppress the host defence response and enable colonisation. The publication of the *P. infestans* T30-4 genome, revealed the presence of 563 effector genes encoding the conserved Arg–any amino acid–Leu–Arg (RXLR) peptide motif (Haas *et al.*, 2009). These effectors, rapidly evolve by gaining and losing repeat-rich domains through recombination with different paralogs, transposon movement, and point mutations (Goss *et al.*, 2013). During co-evolution, potato has evolved receptors to recognise some of these effectors and trigger an immune response.

Wild *Solanum* species are the main source of resistance (*R*) genes to *P. infestans* (*Rpi*). To date, over 20 *Rpi* genes have been characterised in different *Solanum* species, e.g. *R1*, *R2*, *R3a, R3b, R8, R9a* from *S. demissum*, *Rpi-blb1*, 2 from *S. bulbocastanum*, *Rpi-vnt1* from *S. venturii* and *Rpi-amr1* from *S. americanum* (Ballvora *et al.*, 2002; van der Vossen *et al.*, 2003, 2005; Huang *et al.*, 2005; Pel *et al.*, 2009; Lokossou *et al.*, 2009; Li *et al.*, 2011; Jo *et al.*, 2015; Vossen *et al.*, 2016; Witek *et al.*, 2020). All these receptors belong to the nucleotide-binding (NB)–leucine-rich repeat (NLR) type of receptors and contain a coiled-coil domain (CC) in their N-termini, referred to as CC-NB-LRR or CNL. The recognition of a specific effector or a-virulence factor (AVR) leads to the activation of the plant defences and the restriction of the pathogen growth. To keep up with the fast evolution of effectors, NLR genes are also very diverse and rapidly evolving. Gene duplications, recombinations, unequal crossing overs and transpositions have been proposed to provide the basis for the evolution of the NLR recognition spectrum (Leister, 2004; Mcdowell & Simon, 2006). This fast evolution can lead to the independent development of new Rpi receptors in different geographical locations that recognise the same effector. For instance, the recognition of the effector AVR2 from *P. infestans*, by the unrelated R2 and Rpi-mcq1 CNLs (Aguilera-Galvez *et al.*, 2018). R2 is located on chromosome IV in the Mexican species *S. demissum*, while Rpi-mcq1 is located on chromosome IX from a Peruvian accession of *S. mochiquense* (Smilde *et al.*, 2005; Foster *et al.*, 2009). When the doubled-monoploid DM1-3 519 R44 potato genome was published, 755 NLR genes were identified (Jupe *et al.*, 2013). Many of them were found in clusters together with closely related paralogs. All of these clusters were formed in ancestral species and had sequence homology to syntenic genomic regions from other *Solanum* species harbouring late blight resistance genes. Thus, inactive *Rpi* homologs (*rpi*) can be found in all *Solanum* genomes.

Here, we studied *Solanum chacoense* (*S. chacoense*); a diploid wild potato relative from South America considered a source of resistance to *P. infestans*. We identified two functionally distinct receptors, Rpi-chc1.1 and Rpi-chc1.2, which are allelic variants that recognise distinct *P. infestans* effectors from the same PexRD12/31 effector superfamily. Remarkably, only Rpi-chc1.1 is able to provide resistance against current *P. infestans* isolates. The expression and recognition of PexRD12 effectors was associated with Rpi-chc1.1 mediated resistance and, therefore, designated as AVRchc1.1 effectors. PexRD31 effectors were still expressed in current *P. infestans* isolates, but were rapidly downregulated during the interaction with potato. This, potentially explains the inability of *Rpi-chc1.2* to provide late blight resistance. We postulate that *Rpi-chc1.2* is a ubiquitous ancient *R* gene that was recently overcome and PexRD31 may have functioned as AVRchc1.2. An allele mining strategy revealed Rpi-chc1 orthologs in different wild *Solanum* accessions and potato cultivars that could be classified by their sequence and recognition spectrum of AVRchc1.1, AVRchc1.2, or non-functionality. Finally, using domain swaps, we found that the LRR domain harboured the recognition specificity of both AVRchc1.1 and AVRchc1.2. The specificities resided in overlapping LRR subdomains and could not be combined into one active protein using domain exchanges.

## Materials and Methods

### Plant materials and growth conditions

The wild *Solanum* species used in this study are listed in **Table S1** (Tan *et al.*, 2010; Vleeshouwers *et al.*, 2011a). The potato plants were maintained *in vitro* on MS20 at 24°C under 16/8h day/night regime (Domazakis *et al.*, 2017). The 7650 F1 population was generated by crossing *S. chachoense* (CHC543-5) × *S. chacoense* (CHC544-5). *S. tuberosum cv.* ‘Désirée’ was used for stable transformations of the different *Rpi-chc1.1* candidates. Four week old *Nicotiana benthamiana* leaves were used for agroinfiltration. The agroinfiltrated plants were kept in climate regulated greenhouse compartments of Unifarm (Wageningen University & Research) at 20-25°C and under 16/8h day/night regime.

### BAC clone isolation and sequencing

The procedure has been described in patent US9551007B2. Briefly: Two different BAC libraries were produced using partial digestion of CHC543-5 genomic DNA with HindIII. Fragments larger than 100kb were ligated into pBeloBAC or pCC1BAC arms (Epicenter). The BAC clones were collected and stored as bacterial pools of approximatively 700 to 1000 white colonies. BAC pools were screened with selected markers and individual clones were identified using colony PCR. The ends of positive individual BACs were sequenced for the purpose of fine mapping RH106G03T and RH137D14_C37-7-4. The complete inserts were sequenced using shotgun sequencing of 2kb library fragments generated by partial digestion with EcoR1 by Macrogen, Inc (Seoul, South Korea). Assembly of the sequences resulted in contigs as indicated in Fig. **1** (Genbank accession number MW383255).

**Fig. 1.**
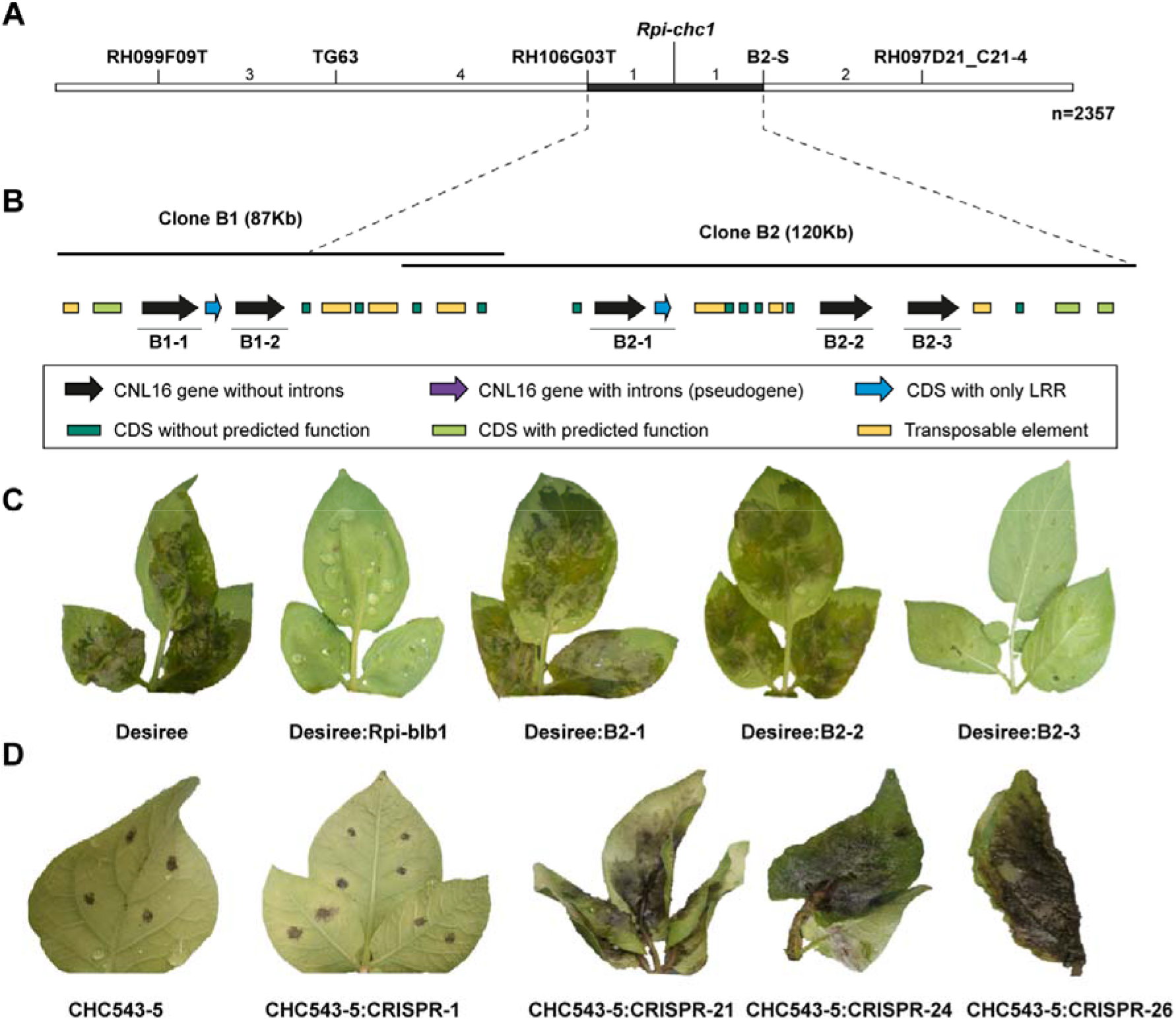
Map based cloning of *Rpi-chc1.1*. (A) Genetic map of *P. infestans* (isolate 90128) resistance from CHC543-5. The number between the markers represents the number of recombinants found in a population derived from 2357 seedlings. Markers starting with RH were derived from BAC end sequences generated by PGSC. Marker B2-S represents the BAC end marker from clone 2. The black horizontal line represents the interval of *Rpi-chc1.1*. (B) Two BAC clones were isolated to generate the physical map. Annotation revealed the presence of NB-LRR genes, genes with or without predicted function and transposable elements. Three complete NB-LRR (B2-1, B2-2 and B2-3) genes between flanking markers RH106G03T and B2-S were selected as candidates. (C) The three candidates were stably transformed into the potato variety Desiree. After inoculation with isolate 90128, only the candidate B2-3 was able to provide resistance. Untransformed Desiree and Desiree plants stably transformed with *Rpi-blb1* were used as negative and positive controls, respectively. (D) CRISPR-Cas9 constructs were designed to specifically target candidate B2-3 and stably transformed in the *S. chacoense* 543-5 resistant genotype. Transgenic plants with B2-3 knock-outs were susceptible to *P. infestans* 90128 and IPO-C isolates. Transgenic plants without mutations in the B2-3 candidate and untransformed CHC543-5 were used as a control.

### Cloning of *Rpi-chc1* allelic variants and chimeric constructs

The *Rpi-chc1* allelic variants were amplified using genomic DNA from the different wild *Solanum* species using PCR primers as described in Table **S2** and DNA polymerase with proofreading activity. The fragments were cloned into pGEM-T easy vector (Promega) for sequencing. Genbank submission numbers as in Table **S1**. The *Rpi-ber1.1* and *Rpi-tar1.1* genes were amplified using primers in the promotor and terminator. The resulting PCR fragments were cloned into pBINPLUS-PASSA (Jo *et al.*, 2016) and were expressed in transgenic Desiree plants under the control of their native regulatory elements. For transient expression analyses, the coding sequences of the allelic variants were cloned under the *Rpi-chc1.1* regulatory elements (900bp promotor and 400bp terminator) into pDEST using a multisite gateway protocol. *Escherichia coli* strain DH10ß was transformed with the gateway reaction products and clones with the correct insert were selected. *Agrobacterium tumefaciens* AGL1+VirG was used for transient and stable transformations of *N. benthamiana* leaves and *S. tuberosum cv.* ′Désirée′.

The chimeric constructs were cloned using the Golden Gate modular cloning principle. As acceptor vector, we used a Golden Gate compatible version of pBINPLUS (McBride & Summerfelt, 1990), PBINPLUS-GG (Vossenberg *et al.*, 2019). The final acceptor vector was constructed to contain 800bp *Rpi-chc1.1* promoter::CDS::1000bp *Rpi-ber* terminator (Fig. **S1**). The different PCR fragments were amplified using the Phusion High-Fidelity PCR Kit (Thermo SCIENTIFIC) and primers with BsaI sites as overhang (Table **S2**) and purified using the DNA Clean&Concentrator Kit (ZYMO RESEARCH). PCR fragments and the acceptor vector were incubated in Buffer G (Thermo SCIENTIFIC) with ATP 1mM for thirty cycles of 37°C for 5min + 16°C for 5min. Additionally, we performed a final step at 37° for 10min, to digest the plasmids wrongly assembled, and 65°C for 20min, to heat inactivate the BsaI enzyme.

### Hypersensitive cell death assays

Transient expression of the different receptors and PexRD12/31 effectors were performed in four weeks old *N. benthamiana* leaves. R3a/AVR3a was used as a positive control. All the constructs were agroinfiltrated at an OD_600_ of 0.5. Each construct was agroinfiltrated twice on two leaves of four plants in at least two independent experiments. Cell death responses were observed after 3-4 days post inoculation.

### Phylogenetic analysis of Rpi-chc1.1 homologs and the PexRD12/31 superfamily

The sequences of the PexRD12/31 effectors were retrieved from the *P. infestans* T30-4 genome (Haas *et al.*, 2009). Twenty family members were found to form the PexRD12/31 superfamily. The coding sequences of the *Rpi-chc1* variants as obtained in this study, were aligned using MUSCLE and a neighbour joining tree was calculated using Megalign from the DNAstar package. The closest homolog of *Rpi-chc1.1* from the DM reference genome (SoltuDM10G021850.1) was used as an outgroup.

The protein sequences of PexRD12/31 effectors were aligned using Clustal OMEGA and manually edited in MEGAX (Sievers *et al.*, 2011; Kumar *et al.*, 2018). The phylogenetic relationship was inferred using the Maximum Likelihood method based on the JTT matrix-based model in MEGAX with 1000 bootstraps (Jones *et al.*, 1992). The tree with the highest log likelihood was shown. The two more distant effectors PITG_16428 and PITG_09577, served as an outgroup.

### *P. infestans* isolates and Detach Leaf Assay (DLA)

The *P. infestans* isolates used in this study (90128, IPO-C and NL08645) were retrieved from our in-house collection. Isolates were grown at 15°C on solid rye medium in the dark (Caten & Jinks, 1968). After two weeks, sporulating mycelium was flooded with 20 mL of ice-cold water, adjusted to 70 zoospores/μL and incubated at 4°C for 2-3 hours. After the incubation, the detached leaves were inoculated with 10μL of the zoospore suspension on the abaxial side of the leaves. Detached leaves were inserted into wet floral foam. For each biological replicate the three leaflets from four leaves from two independent plants were used. Twelve spots on each leaf were inoculated with the zoospore suspension and closed in a plastic bag, to maintain high humidity. The leaves were kept in a climate cell at 18°C for 5 days. Disease resistance was scored on a scale from 1 to 10 for each leaflet. 10=no symptoms; 9=HR no larger than the inoculum droplet; 8=HR lesion of up to 0,5 cm diameter; 7=diffuse lesions up to 1cm diameter, no sporulation, no water soaking; 5= lesions larger than 1 cm sometimes with water soaking, no sporulation; 4=large water soaked lesions with sporulation only visible through binoculars; 2= large lesions with macroscopically visible sporulation on one side of the leaflet; 1= large lesions with macroscopically visible sporulation on both sides of the leaflet.

### Relative effector and *R* gene expression

The *P. infestans* effectors used in this study are listed in Table **S3**. The different genotypes were inoculated with the different *P. infestans* isolates and samples were collected after 0, 3, 8, 24, 48, 72, 96 and 120 hours. Infected plant material with the different *P. infestans* isolates was collected and RNA was isolated using RNA Purification Kit (QIAGEN). The isolated RNA was converted into cDNA using the QuantiTect Reverse Transcription Kit (QIAGEN). The primers used in this study are listed in Table **S2**. The expression of the different effectors in the infected material was evaluated using RT-qPCR SYBR Green (Bio-Rad). The samples were heated to 95°C for 2min. Then 40 cycles of 15sec at 95°C, 30sec at 60°C and 30sec at 72°C. Fluorescence was measured after each cycle. After the final amplification cycle a melting curve was calculated. Relative gene expression was calculated using the 2 ^−ΔΔCT^ method (Livak & Schmittgen, 2001). The normalised gene expression was obtained by dividing the relative gene expression by the relative *P. infestans* elongation factor 2 gene (*ef2*) expression.

### sgRNA and CRISPR-Cas9 construct design

The CRISPOR web tool (http://crispor.org) was used to design the sgRNAs with lower off-target and higher on-target potentials (Concordet & Haeussler, 2018).

A Modular Cloning (MoClo) system based on the Golden Gate cloning technology was used to assemble the different sgRNAs and binary vectors as previously described for tomato mutagenesis (Engler *et al.*, 2008; Weber *et al.*, 2011). Briefly, each sgRNA was fused to the *Arabidopsis thaliana* U6-26 promoter as *AtU6-26::gRNA*. The Level 1 constructs *pICH47732-pNOS::NPTII::tOCS*, *pICH47742-p2×35S::hCas9::tNOS* and the linker *pICH41780* were used to build the Level 2 vector *pICSL4723* (Werner *et al.*, 2012). The primers used for cloning the gRNAs are listed in Table **S2**.

## Results

### Cloning and characterization of *Rpi-chc1.1*

The *S. chacoense* accession CHC543 from Bolivia is a previously described wild potato relative harbouring resistance to *P. infestans* (Vleeshouwers *et al.*, 2011a). To identify the genetic locus of resistance, the resistant seedling CHC543-5 was crossed with the susceptible seedling CHC544-5 to generate the F1 population 7650, consisting initially of 212 individuals. This population was challenged with *P. infestans* isolate 90128 in a detached leaf assay (DLA). A clear 1:1 segregation was observed, indicating the presence of a single dominant resistance gene which will henceforth be referred to as *Rpi-chc1*. CAPS markers from chromosome 10 were tested as this chromosome was known to harbour *Rpi-ber* from the related species *S. berthaultii* (Vossen *et al.*, 2013). The marker TG63 in chromosome 10 was indeed linked to the *Rpi-chc1* resistance. Successive fine mapping in a recombinant population representing 2357 individuals was performed using markers derived from RH89-39-16 BAC clones from chromosome 10 (PGSC) (The Potato Genome Sequencing Consortium, 2011; Sharma *et al.*, 2013). A narrow genetic window between markers RH106G03-T and RH97D21_C21-4 was identified to contain *Rpi-chc1* (Fig. **1a**). To generate a physical map of the mapping interval, two Bacterial Artificial Chromosome (BAC) clones, B1 and B2, were selected from a BAC library that was derived from CHC543-5 genomic DNA. After sequencing the BAC clones, two NLR genes were identified in clone B1 and another six NLR in clone B2. Further fine-mapping revealed that only the last six were located within the mapping interval and only three (B2-1, B2-2 and B2-3) encoded complete NLR proteins (Fig. **1b**). The three candidates were subcloned including their native 5’ and 3’ regulatory elements, and complementation analyses were performed in *N. benthamiana*. After two days, the agroinfiltrated area was challenged with *P. infestans* 90128. *Rpi-blb1*, which was shown to provide resistance to *P. infestans*, was used as a positive control. The leaves agroinfiltrated with candidate B2-3 and *Rpi-blb1* showed severely compromised pathogen growth, while leaves with candidates B2-1 and B2-2 were completely susceptible to *P. infestans* 90128 (Fig. **S2**). This result suggested that B2-3 was the gene in CHC543-5 that provides resistance to *P. infestans*. To verify this result, the three candidates B2-1, B2-2 and B2-3 were stably transformed into the susceptible *S. tuberosum* cv. ‘Désirée’. Indeed, only the events containing candidate B2-3 showed resistance to *P. infestans* (Fig. **1c**). Furthermore, we specifically targeted the B2-3 candidate with sgRNAs and CRISPR-Cas9 enzyme by stable transformation of the resistant CHC543-5 genotype with. The transformation events were challenged with *P. infestans* 90128 and IPO-C isolates, and 48% of the transformants became susceptible to both isolates (Fig. **1d**; Table **S4**). Therefore, we concluded that B2-3 was the gene from CHC543-5 that was causal for late blight resistance. Henceforth, we will refer to gene B2-3 as *Rpi-chc1.1* as it is the first *Rpi-chc1* allele that is identified in *S. chacoense*.

### Identification of *Rpi-chc1.1* allelic variants

In order to identify different *Rpi-chc1.1* allelic variants, we pursued an allele mining approach in several resistant and susceptible *S. chacoense*, *S. berthaultii*, *S. tarijense*, and *S. tuberosum* accessions. Homologous sequences were amplified using primers overlapping the start and stop codons of *Rpi-chc1.1*. The PCR fragments of the expected 3.9 kb size were cloned and sequenced, resulting in the identification of fifteen *Rpi-chc1.1*-like sequences. From the selected diploid accessions one or two sequence variants were identified, suggesting that indeed *Rpi-chc1* alleles were mined rather than paralogs. Phylogenetic analysis of the sequences showed strong sequence similarities among the alleles (94.6 – 100% identity). Even within this high identity range, the presence of four main clades was revealed (Fig. **2a**). In clade 1, the *Rpi-chc1.1* allele was found, together with three sequences from *S. berthaultii* that were (nearly) identical to each other, and a sequence from *S. tarijense*. From clade 1, together with *Rpi-chc1.1*, we selected one sequence from *S. berthaultii* (94-2031) and the *S. tarijense* (TAR852-5) for complementation analysis. Transformation of the corresponding genes to susceptible Désirée plants showed that they provide resistance to *P. infestans* isolates 90128 and IPO-C, like *Rpi-chc1.1* (Table **S5**). We therefore concluded that clade 1 contains functional alleles of *Rpi-chc1*. The *S. tarijense* allele will be referred to as *Rpi-tar1.1*. The *S. berthaultii* allele will be referred to as *Rpi-ber1.1* which matches to the previously described *Rpi-ber* and *Rpi-ber1* genes that were derived from the same accession (PI473331) at similar genetic positions (Rauscher *et al.*, 2006; Tan *et al.*, 2010; Vossen *et al.*, 2013).

**Fig. 2.**
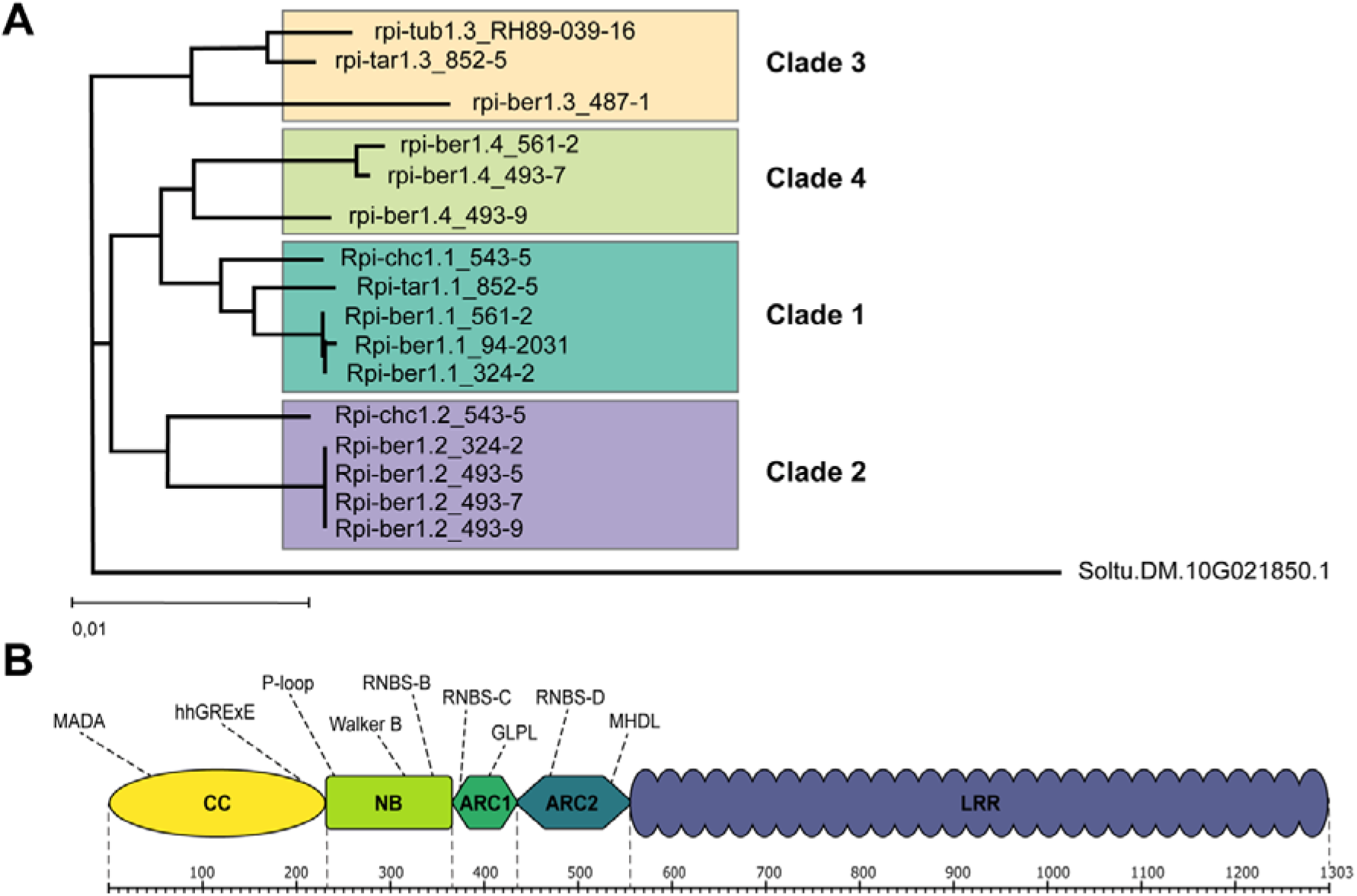
*Rpi-chc1* allele mining. (A) Sixteen *Rpi-chc1.1*-like sequences were cloned from eleven different diploid *Solanum* accessions. From seven accessions two variants were identified. From four accessions only one variant was found, suggesting that the second allele did not match the PCR primers. The phylogenetic analysis of the DNA sequences led to the identification of four clades. The branch lengths represent the percentage of phylogenetic distance. (B) The *Rpi-chc1* alleles belong to the CNL immune receptor family. Different motifs were found in the different receptor domains. The LRR domain consists of 29 imperfect repeats.

The allele mining in accession CHC543-5 resulted not only in the re-identification of the active *Rpi-chc1.1* but also in the identification a presumed allelic variant, which we will refer to as *Rpi-chc1.2*. To test if *Rpi-chc1.1* and *Rpi-chc1.2* were indeed alleles of the same gene, we tested *Rpi-chc1.2* specific markers in the recombinant population 6750 (CHC543-5 × CHC544-5). We found a perfect repulsion between *Rpi-chc1.2* and *Rpi-chc1.1*, strongly suggesting that both genes are allelic variants (Table **S6**). Additionally, this analysis proved that *Rpi-chc1.2* does not cause resistance against *P. infestans* 90128, even though *Rpi-chc1.2* is expressed during infection (Fig. **S3f**). The Rpi-chc1.2 protein sequence clusters in clade 2 together with four identical sequences from *S. berthaultii*. Close to clade 2, we can observe clade 3, which consisted of a *S. berthaultii*, a *S. tarijense* and a *S. tuberosum* allele from RH89-039-16, a diploid clone previously characterised as susceptible to *P. infestans* (Vleeshouwers *et al.*, 2011a). The clade 3 allele from *S. tarijense* contained an in frame stop codon, making it unlikely that this allele is producing an active resistance protein. Additionally, a fourth clade contained only *S. berthaultii* alleles. The allelic variants were numbered according to the clade in which they were found (i.e. *Rpi-ber1.1* from clade 1 and *Rpi-ber1.2* from clade 2, etc.) followed by an extension to indicate the genotype from which the allele was derived.

The mined *Rpi-chc1.1* variants contained between 1296 to 1303 amino acids (Fig. **2b**; Fig. **S4)** and the encoded proteins belong to the CNL-16 immune receptor family (Witek *et al.* 2016). Their CC domains contain the N terminal MADA motif, 4 predicted α-helices and the typical hhGRExE, but the distinctive EDVID motif was less conserved. The NB domain contains the characteristic P-loop or Kinase 1a domain, the VYND motif, Kinase 2 domain or Walker B, and the Kinase 3a or RNBS-B. The ARC1 domain contains the RNBS-C, the Motif 3 and the GLPL motif; and the ARC2 contains the Motif 2, the RNBS-D and two copies of the MHDL motif. The LRR domain consists of 29 imperfect repeats. Both LRR3 and LRR4 contain a central VLDL motif which is conserved in the third LRR of most functional NLRs.

### Rpi-chc1.1 recognises the RXLR PexRD12 effector family from *P. infestans*

To understand the resistance mechanism of the *S. chacoense* CHC543-5 accession, we searched for the effector recognised by Rpi-chc1.1. A collection of ninety *P. infestans* extracellular (Pex) proteins in a PVX agroinfectious vector, of which 54 contained the RXLR-DEER motif (PexRD), was screened. Individual clones from the Pex collection were co-agroinfiltrated with *Rpi-chc1.1* in *N. benthamiana* leaves. As a positive control, we used a mix of the *R3a*/*AVR3a R* gene effector pair which is known to trigger a strong hypersensitive response (HR) in *N. benthamiana* leaves. Only two effectors from the Pex collection were able to trigger an *Rpi-chc1.1* dependent HR, PexRD12-1 and PexRD12-2 (PITG_16233 and PITG_16240, respectively) (Fig. **3a**). Neither the inactive paralogs B2-1 and B2-2 nor R3a produced an HR upon co-agroinfiltration with PexRD12. These results showed that PexRD12 is specifically recognised by Rpi-chc1.1. We could further confirm this finding using transgenic Désirée potato plants that were transformed with *Rpi-chc1.1*. About half of this transgenic population showed late blight resistance while the other half was susceptible, probably due to impaired transgene expression. Interestingly, the plants that showed late blight resistance also showed PexRD12 recognition, while the susceptible transgenic plants did not show any response upon PexRD12 agroinfiltration (Table **S7**). We sought for further evidence that PexRD12 was indeed causing a-virulence on *Rpi-chc1.1* expressing plants. In a field trial with natural infection, we found isolates that were virulent on plants containing *Rpi-chc1.1*. The infected material was collected and used for gene expression analysis via RT-qPCR. The expression of PexRD12 was significantly reduced in the *Rpi-chc1.1* resistance breaking isolates, while other effectors such as *AVRsto1* were normally expressed. Reciprocally, we found that *Rpi-sto1* breaking isolates still expressed PexRD12 normally (Fig. **3b**). Altogether, these results suggest that PexRD12 corresponds to *AVRchc1.1*.

**Fig. 3.**
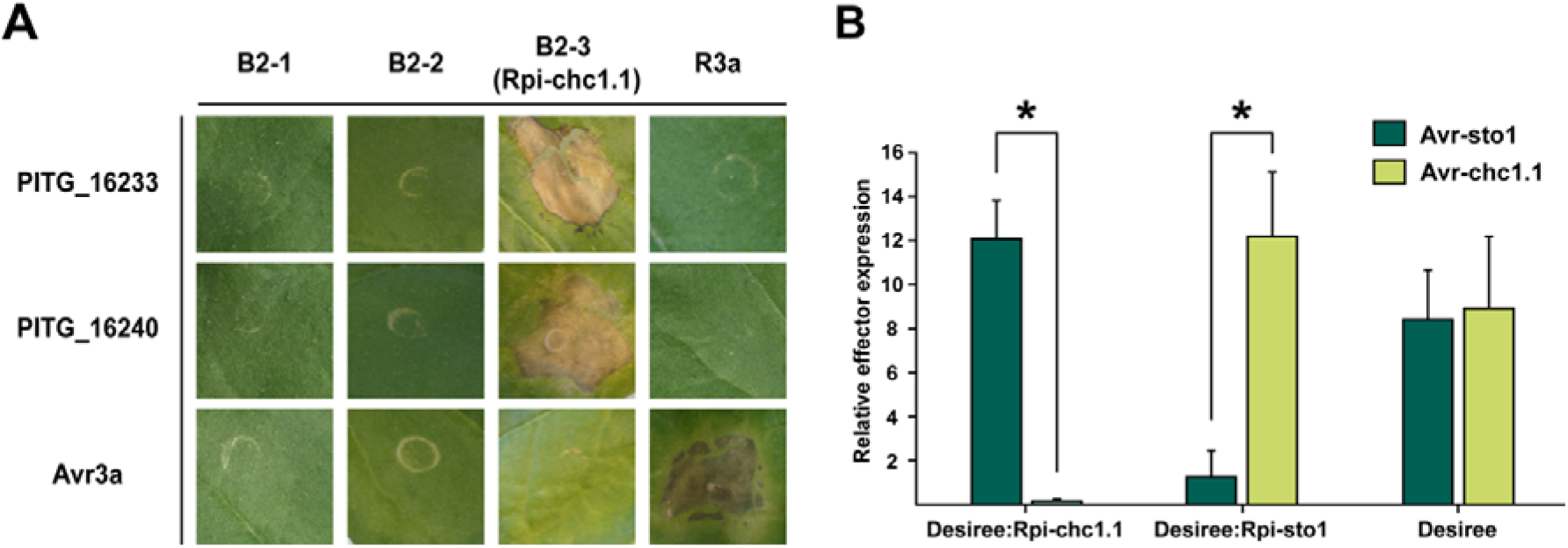
The RXLR effector PexRD12 corresponds to Avrchc1.1. (A) The three *Rpi-chc1.1* candidates were co-infiltrated with the Pex effector collection in *N. benthamiana* leaves to screen for Avrchc1.1. Rpi-chc1.1 induces cell death when co-expressed with both PITG_16233 and PITG_16240, from the PexRD12 family. *R3a* and *Avr3a* were used as negative controls, a mix of *R3a* and *Avr3a* were used as a positive control. (B) Relative *Avrchc1.1* and *Avrsto1* effector expression in *P. infestans* field isolates collected from untransformed Desiree plants, and Desiree plants transformed with Rpi-chc1 (Desiree:*Rpi-chc1.1)*, or Rpi-sto1 (Desiree:*Rpi-sto1*) (2013, Wageningen). Three independent samples were included in the RT-qPCR experiment. Stars represent statistical difference in a two sample T-test, p<0.008.

### The PexRD12/31 superfamily is a complex *P. infestans* RXLR effector family

Using Blast analyses of the T30-4 proteome, we found 9 homologs of PexRD12 in the *P. infestans* T30-4 genome. Additionally, we found that PexRD12 proteins had strong homology with 9 members of the PexRD31 family and two additional, more distantly related sequences (Table **S3**). These 20 effectors are encoded by clusters of paralogs mainly in three supercontigs (Fig. **S5**) and will henceforth be referred to as the PexRD12/31 superfamily (see also Petre *et al.*, 2020). All PexRD12/31 effectors are small proteins that include a signal peptide in the N-terminus, an effector domain in the C-terminus, and the conserved RXLR and EER motifs in the centre; except for PITG_16243 and PITG_09577 which contain an RXXR-EER and RXXLR-EER motifs, respectively (Fig. **4a**).

**Fig. 4.**
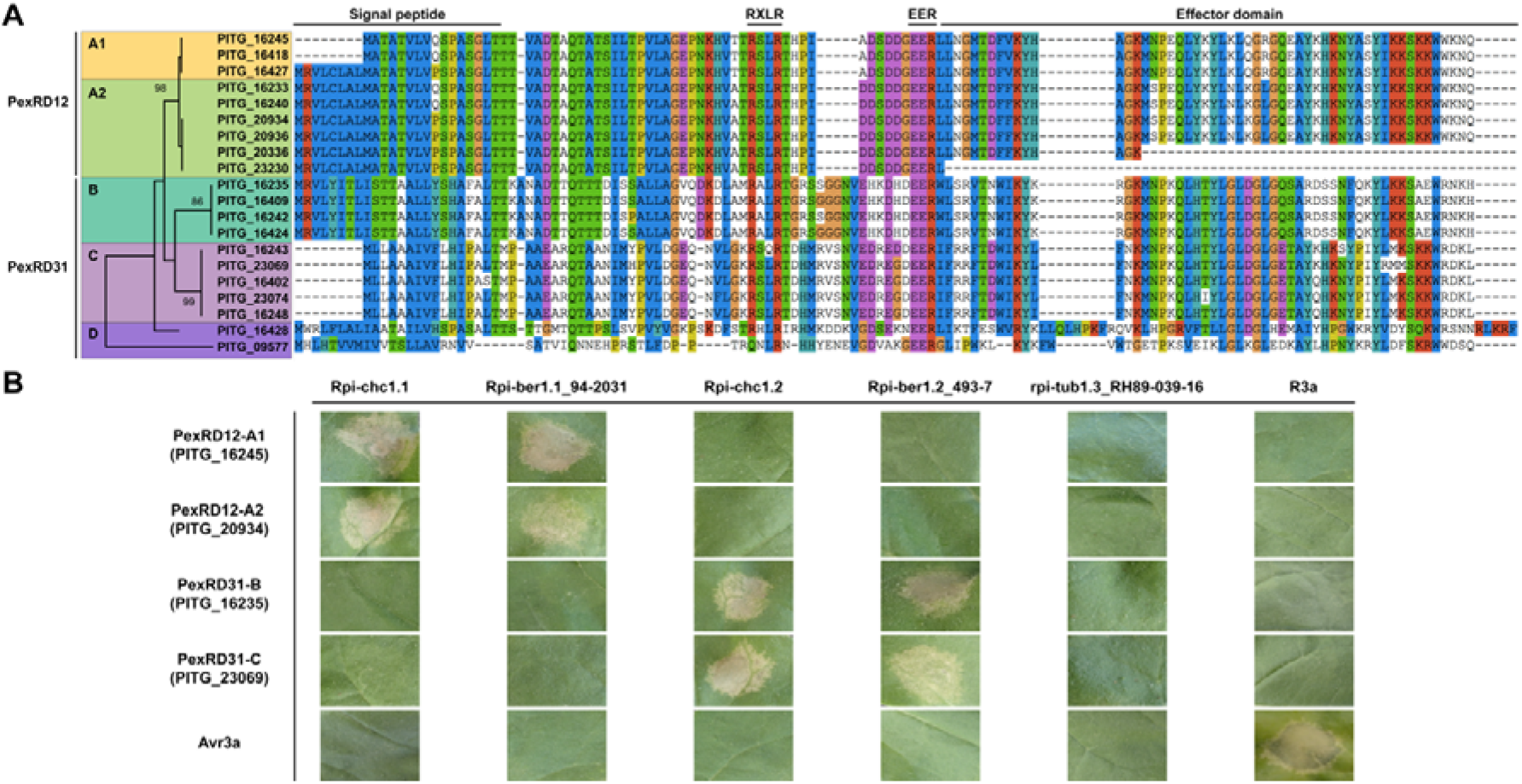
Rpi-chc1 alleles show non-overlapping recognition of the PexRD12/31 effector superfamily. (A) The twenty members of the PexRD12/31 superfamily found in *P. infestans* isolate T30-4. In the amino acid sequence we can distinguish a signal peptide in the N-terminus, the conserved RXLR-EER motifs in the center, and the effector domain in the C-terminus. Some PexRD12/31 family members differed at the nucleotide level but were identical at the protein level (PITG_16245 = PITG_16418; PITG_16233 = PITG_16240; PITG_20934 = PITG_20936; PITG_16409 = PITG_16424). The phylogenetic analysis of the complete protein sequences led to the identification of five clades. This analysis was performed in MEGA X by using the Maximum Likelihood method based on the JTT matrix-based model. The tree with the highest log likelihood (−766) is shown. The bootstrapping values, which indicates the percentage of trees that had the particular branch, are shown in each branch. (B) Different *Rpi-chc1* allelic variants were co-agroinfiltrated in *N. benthamiana* with a member from each PexRD12/31 clade. While variants from clade 1 recognize both PexRD12 A1 and A2 clades, *Rpi-chc1* variants from clade 2 recognize PexRD31 B and C. A mix of *R3a* and *Avr3a* was used as a positive control.

The alignment of the protein sequences and the phylogenetic analysis of the PexRD12/31 superfamily members resulted in five main clades (Fig. **4a**). Two highly homologous clades can be distinguished to form the PexRD12 family, PexRD12-A1 and PexRD12-A2. The clade PexRD12-A2 also includes truncated versions which partly or completely miss the effector domain. In addition, two related clades constitute the PexRD31 family, PexRD31-B and PexRD31-C. Additionally, PITG_16428 and PITG_09577 were much less related and are together referred to as PexRD12/31 group D.

To determine the degree to which PexRD12/31 members are expressed *in planta*, as observed for other AVR effectors of *P. infestans* (Vleeshouwers *et al.*, 2011b; Rietman *et al.*, 2012), we tested their expression during infection with quantitative PCR on cDNA using clade A, B and C specific primers. The relative expression was calculated and normalised for the relative amount of *P. infestans.* Three different *P. infestans* isolates were evaluated at different time points after inoculation of different susceptible potato genotypes (Fig. **S3a-d**). In all the tested genotypes, PexRD12 showed the highest relative expression. In two isolates a maximum expression was found from 4 to 24 hours after inoculation and expression remained high till after 48 hours in all four isolates. The PexRD31-B effectors were expressed in 2 isolates but were rapidly downregulated in the first hours after inoculation with hardly any expression left. The expression of PexRD31-C was mostly undetectable along the inoculation time course.

### Rpi-chc1.2 recognises the RXLR PexRD31 effector family from *P. infestans*

In order to describe the spectrum of effector recognition by different *Rpi-chc1* alleles, several representatives from each clade were selected and co-agroinfiltrated with different PexRD12/31 members in *N. benthamiana*. *Rpi-chc1.1_543-5* and *Rpi-ber1.1_*94-2031-01 from clade 1, *Rpi-chc1.2_543-5* and *Rpi-ber1.2_*493-7 from clade 2, and *rpi-tub1*-RH89-039-16 from clade 3 were selected. As a representation from each of the clades of the PexRD12/31 effector superfamily, we selected: PITG_16245 (PexRD12-A1), PITG_20934 (PexRD12-A2), PITG_16235 (PexRD31-B), and PITG_23069 (PexRD31-C). The different *Rpi-chc1* alleles were co-agroinfiltrated with the PexRD12/31 effectors in *N. benthamiana* leaves. Three days after agroinfiltration, we observed that the members from clade 1, *Rpi-chc1.1* and *Rpi-ber1.1*, specifically recognised both PexRD12-A1 and PexRD12-A2 effectors (Fig. **4b**). This result showed that Rpi-chc1.1 and Rpi-ber1.1 recognise multiple members of the PexRD12 family, suggesting that AVRchc1.1 is encoded by multiple redundant paralogs. On the other hand, Rpi-chc1.2 and Rpi-ber1.2 from clade 2, specifically recognised both PexRD31-B and PexRD31-C effectors (Fig. **4b**). This suggests that multiple PexRD31 paralogs correspond to AVRchc1.2. The selected allele from clade 3, rpi-tub1.3_RH89-039-16, was not able to recognize any of the PexRD12/31 members (Fig. **4b**), showing that clade 3 encodes functionally more distant receptors, and is in agreement with the known susceptibility of RH89-039-16 to *P. infestans* (Vleeshouwers *et al.*, 2008).

### The LRR domain of the Rpi-chc1 variants determines the PexRD12/31 effector recognition specificity

Since the allelic variants of *Rpi-chc1* could be divided into three activity groups, while having an amino acid identity up to 96%, they provide ideal tools to investigate the Rpi-chc1 mechanism of recognition. Therefore, we performed progressive exchanges of the different receptor domains. The chimeric receptors were co-agroinfiltrated with the PexRD12/31 effectors in *N. benthamiana* leaves to evaluate their recognition specificity. First, we selected Rpi-chc1.2 and rpi-tub1.3_RH89-039-16 as representatives of clade B and clade C, respectively. When aligning the protein sequences, 54 single amino acid polymorphisms (SAPs) were found and most of them were located in the LRR domain (Fig. **5a**). As previously mentioned, Rpi-chc1.2 specifically recognises PexRD31-B and PexRD31-C, while rpi-tub1.3_RH89-039-16 does not recognise any of the PexRD12/31 effectors. When the complete rpi-tub1.3_RH89-039-16 LRR domain was exchanged for the Rpi-chc1.2 LRR, the chimeric receptor RH::C2_2-29 was able to recognise both PexRD31-B and PexRD31-C. Reciprocally, the exchange of the Rpi-chc1.2 LRR for the rpi-tub1.3_RH89-039-16 in C2::RH_2-29, led to the inability to recognise any of the PexRD12/31 effectors. This result demonstrates the importance of the LRR domain during the AVRchc1.2 recognition. Additional domain exchanges were performed in order to identify the essential LRR repeats for the effector recognition. The required LRR repeats for the AVRchc1.2 recognition could be narrowed down with the construct RH::C2_14-19 to nine amino acid polymorphisms (Fig**. 5a**). Due to the absence of polymorphisms in the LRR repeats 14 and 15, we can conclude that only LRR repeats from 16 to 19 are required to activate the rpi-tub1.3_RH89-039-16 allele to recognise AVRchc1.2. Interestingly, the majority of these nine amino acid polymorphisms are particularly situated in the solvent exposed domain (xxLxLxxxx) of every LRR repeat. The exchange of any of the solvent exposed residues for the residues present in the inactive allele led to the partial or complete loss of effector recognition (Fig. **5b**).

**Fig. 5.**
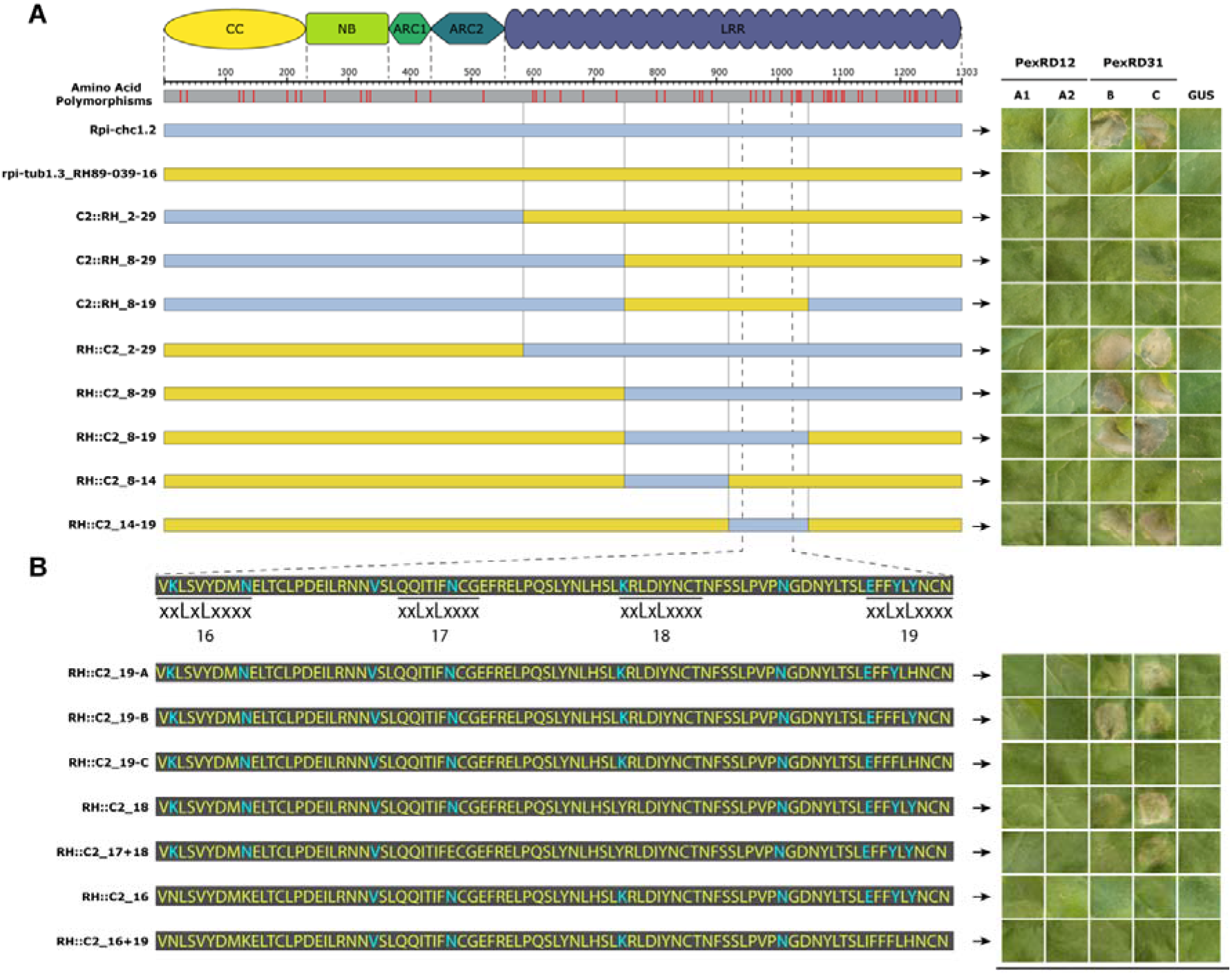
Domain exchanges between rpi-tub1.3_RH89-039-16 and Rpi-chc1.2. (A) The positions of SAPs and the corresponding protein domains are indicated on top. Rpi-chc1.2 and rpi-tub1.3_RH89-039-16, are represented as light blue and yellow bars, respectively. Below, the domain exchanges are shown. The chimeric constructs were co-agroinfiltrated with the PexRD12/31 effectors in *N. benthamiana* leaves. After 4 days, the HR that was visible and recorded. Experiments were repeated three times with 12 inoculation sites each time. A representative leaf for the response of each chimeric construct is shown in the right panel. GUS was used as a negative control. It is concluded that the exchange of the complete LRR domain led to recognition of PexRD31. With the final construct, RH::C2_14-19, the exchange of only nine amino acids led to the activation of the rpi-tub1.3_RH89-036-16 protein. (B) Pinpointing of the amino acids involved in the Rpi-chc1.2 recognition specificity. SAPs are highlighted in blue font. Most of the SAPs are located in the solvent exposed xxLxLxxxx motif of the LRR 16-19. The chimeric constructs were co-agroinfiltrated with the PexRD12/31 effectors in *N. benthamiana* leaves. A representative leaf for the response of each chimeric construct is shown in the right panel. Experiments were repeated three times with 12 inoculation sites each time. GUS was used as a negative control. The modification of any of the Rpi-chc1.2 solvent exposed specific amino acids (blue) for the corresponding amino acid present in rpi-tub1.3_RH89-039-16 (yellow), led to the partial or complete loss of effector recognition.

To understand the difference in effector recognition specificity between Rpi-chc1.1 and Rpi-chc1.2, and to explore the possibility to combine both recognitions in one receptor, we performed a similar progressive domain exchange approach between Rpi-chc1.1 and Rpi-chc1.2 (Fig. **6**). The exchange of the LRR domain in the chimeric receptors C1::C2_8-29 and C2::C1_8-29 led to a shift in effector recognition, from AVRchc1.1 to AVRchc1.2. Further exchanges revealed that the LRR repeats 14 to 23 from Rpi-chc1.2 led to an opposite effector recognition pattern as the chimeric receptor C1::C2_14-23 was able to only recognise AVRchc1.2. With reciprocal domain exchanges of Rpi-chc1.1 into Rpi-chc1.2, we found that LRR repeats 8 to 29, led to AVRchc1.1 recognition. In an attempt to further reduce the length of the exchanged sequence, the recognition of AVRchc1.1 resulted in partial (C2::C1_8-25 and C2::C1_8-23) or complete (C2::C1_14-25 and C2::C1_14-23) loss of recognition. Especially, when comparing the receptors C2::C1_8-29 and C2::C1_8-25, already the modification of the last five SAPs led to the reduced recognition of AVRchc1.1. But, apparently not only the last LRR repeats are involved in the effector recognition. Also the first LRR repeats, from 8 to 14, are also important for AVRchc1.1 recognition as C2::C1_8-25 was able to partially recognise AVRchc1.1, while C2::C1_14-25 did not trigger any HR. We conclude that the LRR repeats 8 to 29 in Rpi-chc1.1 are important for the AVRchc1.1 recognition, which overlaps with LRR repeats 16-19 from Rpi-chc1.2 which were required for AVRchc1.2 recognition.

**Fig. 6.**
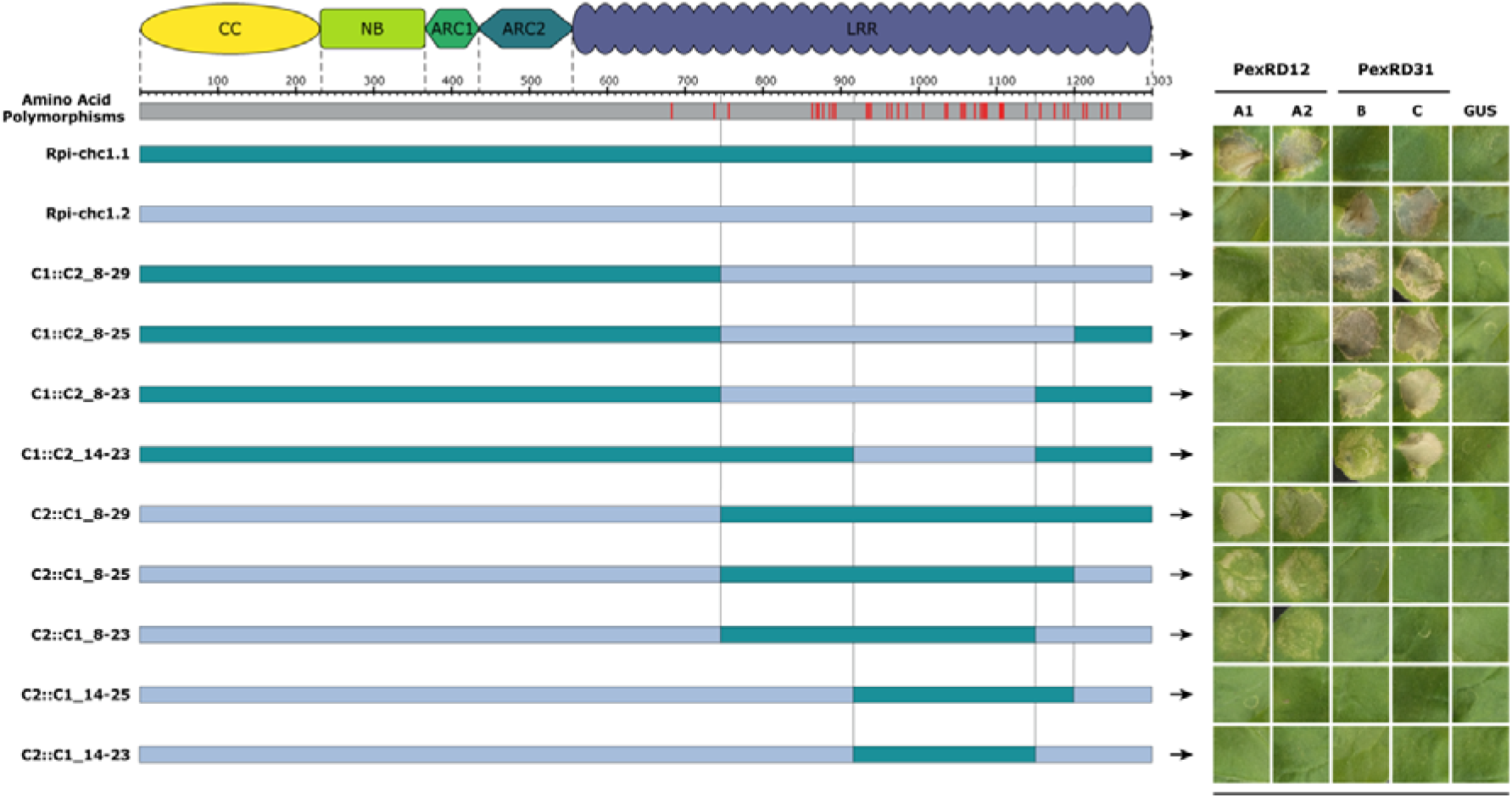
The effector recognition specificity could be exchanged between Rpi-chc1.1 and Rpi-chc1.2. The alignment of Rpi-chc1.1 and Rpi-chc1.2 shows that all the 41 amino acids polymorphisms (red bars) are located in the LRR domain. The chimeric constructs were co-agroinfiltrated with the PexRD12/31 effectors in *N. benthamiana* leaves. A representative leaf for the response of each chimeric construct is shown in the right panel. Experiments were repeated three times with 12 inoculation sites each time. GUS was used as a negative control. In the construct C1::C2_14-23, we could see that the LRR 16-19 reappear again as determining for the PexRD31 recognition. The required domain exchanges of the Rpi-chc1.1 LRR are more complex and encompasses almost the complete LRR.

## Discussion

In this study, we identified *Rpi-chc1.1* and 15 additional allelic variants from *S. berthaultii*, *S. tarijense* and *S. tuberosum*. Phylogenetic analysis of the encoded protein sequences revealed four clades. These four clades were not only supported by sequence similarity but also by differences in effector and *P. infestans* recognition. Clade 1 genes encode receptors that recognise PexRD12 effectors and includes the active orthologs *Rpi-chc1.1*, *Rpi-ber1.1* and *Rpi-tar1.1* (Fig. **2**, **3**). Clade 2 receptors could be distinguished by the recognition of the PexRD31 effectors (Fig. **4**). Receptors encoded by clades 3 and 4 do not recognise PexRD12/31 effectors and no other activity has been found. Interestingly, clade 3 alleles are also present in domesticated potato clones that are susceptible to late blight; e.g. RH89-039-16 (Fig. **2**) and the varieties Colomba and Altus (unpublished data), implying that the encoded receptors are not able to effectively provide resistance against *P. infestans*.

*Rpi-ber1.1_94-2031-01* was derived from the same accession as the previously described *Rpi-ber* (Rauscher *et al.*, 2006; Vossen *et al.*, 2009; Tan *et al.*, 2010), and *Rpi-ber1* genes (Park *et al.*, 2009). In these four studies, *Rpi-ber* and *Rpi-ber1* mapped close to marker TG63 but slightly different genetic positions were reported.

The population from Park *et al*. was quite small and a single recombination event may have caused the deviating genetic distance. In the case of Tan *et al.*, a single mis-phenotyping could explain the mapping of *Rpi-ber* distal to TG63. We therefore assume that *Rpi-ber* and *Rpi-ber1* are the same genes and adopt the *Rpi-ber1* naming as it is more consistent with current nomenclature for late blight resistance genes. *Rpi-ber2,* as described by Park *et al*., was derived from the same accession that was used in our allele mining studies (BER493). We could not find a clade 1 *Rpi-chc1* allele from the BER493 accessions, which supports the idea that a more distantly related CNL16 member may be present that lacks sufficient match to the primer sequences, explaining the *Rpi-ber2* map position distal from TG63.

The presence of *Rpi-chc1* alleles in *S. tarijense* and *S. berthaultii* suggests a functional common ancestor existed before their speciation. However, it must be noted that the geographic locations where the accessions were found are close to each other in Bolivia. Since *S. chacoense*, *S. tarijense* and *S. berthaultii* are closely related, the presence of functional *Rpi-chc1* alleles in the three species might be a result from a recent species intercrossing.

Sequence similarity among the studied allelic variants correlated with their functionality, deduced by their ability to provide late blight resistance and *P. infestans* effector recognition (Fig. **3**, **4**). This is not the first described case of *R* gene allelic variants across *Solanum* species. *Rpi-blb1*, *Rpi-sto1* and *Rpi-pta1*, from the Mexican species *S. bulbocastanum*, *S. stoloniferum* and *S. papita* are allelic variants that recognise the same IpiO or PexRD6 *P. infestans* effector (Vleeshouwers *et al.*, 2008). Among allelic variants of late blight resistance genes (i.e. Rpi-blb3 and Rpi-hjt1 that recognise AVR2 effectors), overlapping recognition specificities have been previously described (Champouret, 2010). Moreover, highly similar, but non-allelic *R* genes from the same CNL cluster had different recognition specificities, i.e. Rpi-vnt1, Rpi-mcq1, R9a, Ph-3 (Smilde *et al.*, 2005; Foster *et al.*, 2009; Zhang *et al.*, 2014; Jo *et al.*, 2015). In the current report, we describe for the first time that allelic variants of a late blight resistance gene show non-overlapping effector recognition specificities. Remarkably, the recognized effectors belonged to the same effector family, which is an intriguing finding in the light of host pathogen co-evolution.

When studying Rpi-chc1 protein domain structure, we identified most of the conserved CNL motifs. Remarkably, the MADA motif (Adachi *et al.*, 2019) was not located downstream of the starting methionine, but downstream of the second methionine in position 46 of the Rpi-chc1 protein. Further research is needed to show if either or both methionines are used as translational start codons. Interestingly, we recently cloned the functional late blight resistance gene from the late blight resistant variety Carolus (*Rpi-Carolus* gene; Hamed Salehian, unpublished data). Rpi-Carolus differed only at 7 amino acid positions from Rpi-ber1, but its N-terminus was shorter as a stop codon was present between the first two methionine codons. This strongly suggests that translation in Rpi-Carolus starts from the second methionine while retaining biological activity.

In contrast to the relatively conserved N termini of the proteins encoded by the *Rpi-chc1* alleles, most interallelic sequence variation localised to the Leucine Rich Repeat regions. Through domain interchange between the *rpi-tub1.3_RH89-039-16* and a *Rpi-chc1.2* alleles and between Rpi-chc1.1 and Rpi-chc1.2, we discovered that the LRR domain defines recognition specificity (Fig. **5**, **6**). Polymorphisms in the LRR of some NLR receptors were previously shown to determine the effector recognition specificity (Dodds *et al.*, 2001; Shen *et al.*, 2003; Catanzariti *et al.*, 2010; Krasileva *et al.*, 2010; Ravensdale *et al.*, 2012; Lindner *et al.* 2020). In one example, a domain exchange between Rx1 and Gpa2 converted the virus resistance into nematode resistance, and vice versa (Slootweg *et al.*, 2017). The recognition of both nematode and virus could not be combined into one chimeric receptor, as we also observed with Rpi-chc1.1 and Rpi-chc1.2. The reason for this is the overlap between the LRRs involved in recognition.

Most of the amino acids in Rpi-chc1.2 that were required for AVRchc1.2 recognition are located in the LRR solvent exposed motif (xxLxLxxxx) and modification of any of the solvent exposed amino acids led to the partial or complete loss of PexRD31 recognition (Fig. **5b**). The co-requirement of these solvent exposed amino acids suggests that they are involved in recognition of a particular epitope. This observation, combined with the observation of unequal distribution of SAPs, allow us to hypothesise that *Rpi-chc1* alleles evolved through insertion of a stretch of DNA into the LRR domain rather than through accumulation of independent mutation. A similar model of evolution was recently proposed for allelic variants of *Rpi-amr1* (Witek *et al.*, 2020). Such insertions may happen through unequal crossing-over with paralog sequences or through retro-transposition. Interestingly, the evolution of integrated domains in *R* genes has been postulated to be caused by an unknown recombination or transposon independent translocation mechanism (Bailey *et al.*, 2018). The same mechanism may be active in LRR exchange to evolve recognition of non-integrated domains like guardees or decoys (Kourelis & van der Hoorn, 2018) or direct effector recognition.

Interestingly, some of the PexRD31 family members have been previously identified as one of the most rapidly diversifying and fast evolving RXLR effectors in the T30-4 genome, with ω values higher than 1.55 (Haas *et al.*, 2009). Additionally, several members of the PexRD12/31 superfamily have recently been characterised to target the host vesicle trafficking machinery by interacting with the vesicle associated membrane protein 72 (VAMP72) family (Petre *et al.*, 2020). Even though both PexRD12 and PexRD31 have the same or functionally similar host targets, they are differentially expressed during *P. infestans* infection. While PexRD12 is highly expressed in all the tested isolates, PexRD31 is expressed at low levels after contact with potato (Fig. **S3**). This would also explain why *Rpi-chc1.2* is not able to provide resistance against *P. infestans*, since most of the isolates have low expression levels of *AVRchc1.2*. Consequently, clade A (PexRD12) may have evolved to avoid detection by *Rpi-chc1.2* while retaining its targeting of the vesicle trafficking machinery.

Another step in the co-evolution between *Rpi-chc1.1* and the PexRD12/31 family was found by analysing the effector expression in plants expressing *Rpi-chc1.1*. The isolates that overcome the *Rpi-chc1.1* resistance no longer express PexRD12, while the expression in untransformed Désirée plants was normal and comparable to the expression of *AVRsto1* (Fig. **3b**). Similarly, evasion of recognition through transcriptional suppression, was previously observed in plants expressing *Rpi-vnt1* infected with *P. infestans* (Pel, 2010). Once more, we confirmed the plasticity of the *P. infestans* effector secretion and the fast evolution capacity of some aggressive isolates to break down single Rpi resistances.

The introgression of single *R* genes is driving *P. infestans* to evolve and evade recognition. In order to durably deploy late blight resistance in agriculture, we need novel strategies informed by knowledge of disease resistance genes in varieties, their recognition specificities and the presence of the cognate effectors in the pathogen populations. Virulence information from the field must be rapidly translated to decision support systems (DSS) for the risk prediction and calculation of biocide spraying intervals. Additionally, DSS can be used to determine *R* gene composition of (novel) varieties to be deployed in the next season. To meet these requirements, novel breeding strategies are needed to rapidly tailor the *R* gene contents of the potato varieties to the pathogen populations. In current breeding schemes it takes 10-15 years to select a late blight resistant potato variety. Moreover, susceptible varieties with dominant market shares will not be easy to replace. A system of varieties with flexible *R* gene content is needed. In other crops this has been accomplished through F1 hybrid varieties. In potato, this route has only recently been opened (Su *et al.*, 2020) and no hybrid potato varieties have reached the market yet. Proof of principle for flexible late blight resistance varieties produced through cisgenesis was provided several years ago (Haverkort *et al.*, 2016). Unfortunately, the EU legislation does not distinguish between cisgenic and transgenic products, making it impossible to market cisgenic varieties. Now, novel gene editing tools have become available, and legislation for their application in agriculture is still under debate. Knowledge as obtained in this study is essential to pursue such applications. We now know how inactive resistance genes from susceptible varieties could be repaired by replacing minimal fragments with the corresponding fragments of alleles from wild relatives. This would provide an unprecedented accuracy and speed which is not in introgression breeding.

## Supporting information

Supplementary data Rpi-chc1 manuscript DM, JV

## Acknowledgements

The first part of this research was performed in the DuRPh project, funded by the Ministry of Agriculture, Nature and Food Quality in the Netherlands. The second part of the research was funded by the Ministry of Infrastructure and Water Management in the Netherlands through the NWO-TTW program Biotechnology and Safety (projectnumber 15815). We thank Jan de Boer and Adillah Tan for advice in selecting BAC clones and marker sequences from chr10. Evert Jacobsen and Clemens van der Wiel are thanked for their advice about regulation and safety aspects of plant biotechnology. Sidrat Abdullah is thanked for testing late blight resistance of transgenic potato plants harbouring *Rpi-chc1* allelic variants. Last but not least, we thank Marjan Bergervoet, Gert van Arkel, Koen Pelgrom, Dirk Jan Huigen and Isolde Pereira for plant transformations, molecular biology assistance and plant maintenance.

## Data Availability

All described sequences have been submitted top GenBank.

## Author contribution

JV planned and designed the research; DML, MN and LK performed the majority of the experiments; SK provided the Pex-RD set; DML and JV wrote the manuscript; RV proofread the manuscript and provided the essential research environment. SA, HS and KS contributed by mapping, cloning and characterization of *Rpi-chc1* allelic variants. RS, AL and AAH contributed through the identification of AVRchc1 and their differential recognition specificities by *Rpi-chc1* allelic variants.

## Supporting Information

Fig. S1: pBINPLUS-PASSA-GG vector map.

Fig. S2: Inoculation of *P. infestans* on *N. benthamiana* leaves agroinfiltrated with *Rpi-chc1.1* candidates.

Fig. S3: Effector and R gene expression in potato leaves inoculated with P infestans.

Fig. S4: Rpi-chc1.1 protein domain organization.

Fig. S5: Localization of PexRD12/31 effectors in the *P. infestans* T30-4 contigs.

Table S1: Accession numbers of Solanum genotypes and Rpi-chc1 sequences

Table S2: Primers used in this study.

Table S3: *P. infestans* effectors used in this study.

Table S4: CRISPR-Cas9 targeting of Rpi-chc1.1.

Table S5: Late blight resistance assessment of different Rpi-chc1 alleles.

Table S6: Segregation of markers and late blight resistance of Rpi-chc1.1 and Rpi-chc1.2.

Table S7: Functional expression of Rpi-chc1.1 in Desiree transgenic events correlates with responsiveness to PexRD12.

## References

Adachi H, Contreras MP, Harant A, Wu C, Derevnina L, Sakai T, Duggan C, Moratto E, Bozkurt TO, Maqbool A, et al. 2019. An N-terminal motif in NLR immune receptors is functionally conserved across distantly related plant species. eLife 8: e49956.

Aguilera-Galvez C, Champouret N, Rietman H, Lin X, Wouters D, Chu Z, Jones JDG, Vossen JH, Visser RGF, Wolters PJ, et al. 2018. Two different R gene loci co-evolved with Avr2 of Phytophthora infestans and confer distinct resistance specificities in potato. Studies in Mycology 89: 105–115.

Bailey PC, Schudoma C, Jackson W, Baggs E, Dagdas G, Haerty W, Moscou M, Krasileva KV. 2018. Dominant integration locus drives continuous diversification of plant immune receptors with exogenous domain fusions. Genome Biology 19: 23.

Ballvora A, Ercolano MR, Weiss J, Meksem K, Bormann CA, Oberhagemann P, Salamini F, Gebhardt C. 2002. The R1 gene for potato resistance to late blight (Phytophthora infestans) belongs to the leucine zipper/NBS/LRR class of plant resistance genes. The Plant Journal 30: 361–371.

Callaway E. 2013. Pathogen genome tracks Irish potato famine back to its roots. Nature: nature.2013.13021.

Catanzariti A-M, Dodds PN, Ve T, Kobe B, Ellis JG, Staskawicz BJ. 2010. The AvrM Effector from Flax Rust Has a Structured C-Terminal Domain and Interacts Directly with the M Resistance Protein. Molecular Plant-Microbe Interactions® 23: 49–57.

Caten CE, Jinks JL. 1968. Spontaneous variability of single isolates of Phytophthora infestans. I. Cultural variation. Canadian Journal of Botany 46: 329–348.

Champouret N. 2010. Functional genomics of Phytophthora infestans effectors and Solanum resistance genes. PhD thesis, Wageningen University. https://edepot.wur.nl/138174

Concordet J-P, Haeussler M. 2018. CRISPOR: intuitive guide selection for CRISPR/Cas9 genome editing experiments and screens. Nucleic Acids Research 46: W242–W245.

Dodds PN, Lawrence GJ, Ellis JG. 2001. Six Amino Acid Changes Confined to the Leucine-Rich Repeat β-Strand/β-Turn Motif Determine the Difference between the P and P2 Rust Resistance Specificities in Flax. The Plant Cell 13: 163–178.

Domazakis E, Lin X, Aguilera-Galvez C, Wouters D, Bijsterbosch G, Wolters PJ, Vleeshouwers VGAA. 2017. Effectoromics-Based Identification of Cell Surface Receptors in Potato. In: Shan L, He P, eds. Methods in Molecular Biology. Plant Pattern Recognition Receptors. New York, NY: Springer New York, 337–353.

Engler C, Kandzia R, Marillonnet S. 2008. A One Pot, One Step, Precision Cloning Method with High Throughput Capability (HA El-Shemy, Ed.). PLoS ONE 3: e3647.

Foster SJ, Park T-H, Pel M, Brigneti G, Śliwka J, Jagger L, van der Vossen E, Jones JDG. 2009. Rpi-vnt1.1, a Tm-2 Homolog from Solanum venturii, Confers Resistance to Potato Late Blight. Molecular Plant-Microbe Interactions® 22: 589–600.

Fry W. 2008. Phytophthora infestans[: the plant (and R gene) destroyer. Molecular Plant Pathology 9: 385–402.

Goss EM, Press CM, Grünwald NJ. 2013. Evolution of RXLR-Class Effectors in the Oomycete Plant Pathogen Phytophthora ramorum (A Palsson, Ed.). PLoS ONE 8: e79347.

Haas BJ, Kamoun S, Zody MC, Jiang RHY, Handsaker RE, Cano LM, Grabherr M, Kodira CD, Raffaele S, Torto-Alalibo T, et al. 2009. Genome sequence and analysis of the Irish potato famine pathogen Phytophthora infestans. Nature 461: 393–398.

Haverkort AJ, Boonekamp PM, Hutten R, Jacobsen E, Lotz LAP, Kessel GJT, Vossen JH, Visser RGF. 2016. Durable Late Blight Resistance in Potato Through Dynamic Varieties Obtained by Cisgenesis: Scientific and Societal Advances in the DuRPh Project. Potato Research 59: 35–66.

Huang S, Van Der Vossen EAG, Kuang H, Vleeshouwers VGAA, Zhang N, Borm TJA, Van Eck HJ, Baker B, Jacobsen E, Visser RGF. 2005. Comparative genomics enabled the isolation of the R3a late blight resistance gene in potato: Cloning the potato late blight R3a gene by synteny. The Plant Journal 42: 251–261.

Huang S, Vleeshouwers VGAA, Werij JS, Hutten RCB, van Eck HJ, Visser RGF, Jacobsen E. 2004. The R3 Resistance to Phytophthora infestans in Potato is Conferred by Two Closely Linked R Genes with Distinct Specificities. Molecular Plant-Microbe Interactions® 17: 428–435.

Jo K-R, Visser RGF, Jacobsen E, Vossen JH. 2015. Characterisation of the late blight resistance in potato differential MaR9 reveals a qualitative resistance gene, R9a, residing in a cluster of Tm-2 2 homologs on chromosome IX. Theoretical and Applied Genetics 128: 931–941.

Jo K-R, Zhu S, Bai Y, Hutten RCB, Kessel GJT, Vleeshouwers VGAA, Jacobsen E, Visser RGF, Vossen JH. 2016. Problematic Crops: 1. Potatoes: Towards Sustainable Potato Late Blight Resistance by Cisgenic *R* Gene Pyramiding. In: Collinge DB, ed. Plant Pathogen Resistance Biotechnology. Hoboken, NJ: John Wiley & Sons, Inc, 171–191.

Jones DT, Taylor WR, Thornton JM. 1992. The rapid generation of mutation data matrices from protein sequences. Bioinformatics 8: 275–282.

Jupe F, Witek K, Verweij W, Śliwka J, Pritchard L, Etherington GJ, Maclean D, Cock PJ, Leggett RM, Bryan GJ, et al. 2013. Resistance gene enrichment sequencing (RenSeq) enables reannotation of the NB-LRR gene family from sequenced plant genomes and rapid mapping of resistance loci in segregating populations. The Plant Journal 76: 530–544.

Kourelis J, van der Hoorn RAL. 2018. Defended to the Nines: 25 Years of Resistance Gene Cloning Identifies Nine Mechanisms for R Protein Function. The Plant Cell 30: 285–299.

Krasileva KV, Dahlbeck D, Staskawicz BJ. 2010. Activation of an Arabidopsis Resistance Protein Is Specified by the in Planta Association of Its Leucine-Rich Repeat Domain with the Cognate Oomycete Effector. The Plant Cell 22: 2444–2458.

Kumar S, Stecher G, Li M, Knyaz C, Tamura K. 2018. MEGA X: Molecular Evolutionary Genetics Analysis across Computing Platforms (FU Battistuzzi, Ed.). Molecular Biology and Evolution 35: 1547–1549.

Leister D. 2004. Tandem and segmental gene duplication and recombination in the evolution of plant disease resistance genes. Trends in Genetics 20: 116–122.

Li G, Huang S, Guo X, Li Y, Yang Y, Guo Z, Kuang H, Rietman H, Bergervoet M, Vleeshouwers VGGA, et al. 2011. Cloning and Characterization of R3b; Members of the R3 Superfamily of Late Blight Resistance Genes Show Sequence and Functional Divergence. Molecular Plant-Microbe Interactions® 24: 1132–1142.

Lindner S, Keller B, Singh SP, Hasenkamp Z, Jung E, Müller MC, Bourras S and Keller B 2020. Single residues in the LRR domain of the wheat PM3A immune receptor can control the strength and the spectrum of the immune response. Plant Journal 104: 200–214.

Livak KJ, Schmittgen TD. 2001. Analysis of Relative Gene Expression Data Using Real-Time Quantitative PCR and the 2−ΔΔCT Method. Methods 25: 402–408.

Lokossou AA, Park T, van Arkel G, Arens M, Ruyter-Spira C, Morales J, Whisson SC, Birch PRJ, Visser RGF, Jacobsen E, et al. 2009. Exploiting Knowledge of R/Avr Genes to Rapidly Clone a New LZ-NBS-LRR Family of Late Blight Resistance Genes from Potato Linkage Group IV. Molecular Plant-Microbe Interactions® 22: 630–641.

McBride KE, Summerfelt KR. 1990. Improved binary vectors for Agrobacterium-mediated plant transformation. Plant Molecular Biology 14: 269–276.

Mcdowell JM, Simon SA. 2006. Recent insights into R gene evolution. Molecular Plant Pathology 7: 437–448.

Park T-H, Foster S, Brigneti G, Jones JDG. 2009. Two distinct potato late blight resistance genes from Solanum berthaultii are located on chromosome 10. Euphytica 165: 269–278.

Pel MA. 2010. Mapping, isolation and characterization of genes responsible for late blight resistance in potato. PhD thesis, Wageningen University. https://edepot.wur.nl/138132

Pel MA, Foster SJ, Park T-H, Rietman H, van Arkel G, Jones JDG, Van Eck HJ, Jacobsen E, Visser RGF, Van der Vossen EAG. 2009. Mapping and Cloning of Late Blight Resistance Genes from *Solanum venturii* Using an Interspecific Candidate Gene Approach. Molecular Plant-Microbe Interactions® 22: 601–615.

Petre B, Contreras MP, Bozkurt TO, Schattat MH, Sklenar J, Schornack S, Abd-El-Haliem A, Castells-Graells R, Lozano-Duran R, Dagdas YF, et al. 2020. Host-interactor screens of Phytophthora infestans RXLR proteins reveal vesicle trafficking as a major effector-targeted process. Plant Biology.

Rauscher GM, Smart CD, Simko I, Bonierbale M, Mayton H, Greenland A, Fry WE. 2006. Characterization and mapping of Rpi-ber, a novel potato late blight resistance gene from Solanum berthaultii. Theoretical and Applied Genetics 112: 674–687.

Ravensdale M, Bernoux M, Ve T, Kobe B, Thrall PH, Ellis JG, Dodds PN. 2012. Intramolecular Interaction Influences Binding of the Flax L5 and L6 Resistance Proteins to their AvrL567 Ligands (J-R Xu, Ed.). PLoS Pathogens 8: e1003004.

Rietman H, Bijsterbosch G, Cano LM, Lee H-R, Vossen JH, Jacobsen E, Visser RGF, Kamoun S, Vleeshouwers VGAA. 2012. Qualitative and Quantitative Late Blight Resistance in the Potato Cultivar Sarpo Mira Is Determined by the Perception of Five Distinct RXLR Effectors. Molecular Plant-Microbe Interactions® 25: 910–919.

Sharma SK, Bolser D, de Boer J, Sønderkær M, Amoros W, Carboni MF, D’Ambrosio JM, de la Cruz G, Di Genova A, Douches DS, et al. 2013. Construction of Reference Chromosome-Scale Pseudomolecules for Potato: Integrating the Potato Genome with Genetic and Physical Maps. G3: Genes|Genomes|Genetics 3: 2031–2047.

Shen Q-H, Zhou F, Bieri S, Haizel T, Shirasu K, Schulze-Lefert P. 2003. Recognition Specificity and RAR1/SGT1 Dependence in Barley Mla Disease Resistance Genes to the Powdery Mildew Fungus. The Plant Cell 15: 732–744.

Sievers F, Wilm A, Dineen D, Gibson TJ, Karplus K, Li W, Lopez R, McWilliam H, Remmert M, Söding J, et al. 2011. Fast, scalable generation of high[quality protein multiple sequence alignments using Clustal Omega. Molecular Systems Biology 7: 539.

Slootweg E, Koropacka K, Roosien J, Dees R, Overmars H, Lankhorst RK, van Schaik C, Pomp R, Bouwman L, Helder J, et al. 2017. Sequence Exchange between Homologous NB-LRR Genes Converts Virus Resistance into Nematode Resistance, and Vice Versa. Plant Physiology 175: 498–510.

Smilde WD, Brigneti G, Jagger L, Perkins S, Jones JDG. 2005. Solanum mochiquense chromosome IX carries a novel late blight resistance gene Rpi-moc1. Theoretical and Applied Genetics 110: 252–258.

Song J, Bradeen JM, Naess SK, Raasch JA, Wielgus SM, Haberlach GT, Liu J, Kuang H, Austin-Phillips S, Buell CR, et al. 2003. Gene RB cloned from Solanum bulbocastanum confers broad spectrum resistance to potato late blight. Proceedings of the National Academy of Sciences 100: 9128–9133.

Su Y, Viquez-Zamora M, den Uil D, Sinnige J, Kruyt H, Vossen J, Lindhout P, van Heusden S. 2020. Introgression of Genes for Resistance against Phytophthora infestans in Diploid Potato. American Journal of Potato Research 97: 33–42.

Tan MYA, Hutten RCB, Visser RGF, van Eck HJ. 2010. The effect of pyramiding Phytophthora infestans resistance genes R Pi-mcd1 and R Pi-ber in potato. Theoretical and Applied Genetics 121: 117–125.

The Potato Genome Sequencing Consortium. 2011. Genome sequence and analysis of the tuber crop potato. Nature 475: 189–195.

Vleeshouwers VG, Finkers R, Budding D, Visser M, Jacobs MM, van Berloo R, Pel M, Champouret N, Bakker E, Krenek P, et al. 2011a. SolRgene: an online database to explore disease resistance genes in tuber-bearing Solanum species. BMC Plant Biology 11: 116.

Vleeshouwers VGAA, Raffaele S, Vossen JH, Champouret N, Oliva R, Segretin ME, Rietman H, Cano LM, Lokossou A, Kessel G, et al. 2011b. Understanding and Exploiting Late Blight Resistance in the Age of Effectors. Annual Review of Phytopathology 49: 507–531.

Vleeshouwers VGAA, Rietman H, Krenek P, Champouret N, Young C, Oh S-K, Wang M, Bouwmeester K, Vosman B, Visser RGF, et al. 2008. Effector Genomics Accelerates Discovery and Functional Profiling of Potato Disease Resistance and Phytophthora Infestans Avirulence Genes (HA El-Shemy, Ed.). PLoS ONE 3: e2875.

Vossen JH, van Arkel G, Bergervoet M, Jo K-R, Jacobsen E, Visser RGF. 2016. The Solanum demissum R8 late blight resistance gene is an Sw-5 homologue that has been deployed worldwide in late blight resistant varieties. Theoretical and Applied Genetics 129: 1785–1796.

Vossen JH, Dezhsetan S, Esselink D, Arens M, Sanz MJ, Verweij W, Verzaux E, van der Linden C. 2013. Novel applications of motif-directed profiling to identify disease resistance genes in plants. Plant Methods 9: 37.

Vossen JH, Nijenhuis M, Arens M, Van Der Vossen EAG, Jacobsen E, Visser RGF. 2009. Cloning and exploitation of a functional R gene from Solanum chacoense. Patent application WO2011034433 A1. Published by the world intellectual property organization 18 September 2009

van der Vossen EAG, Gros J, Sikkema A, Muskens M, Wouters D, Wolters P, Pereira A, Allefs S. 2005. The Rpi-blb2 gene from Solanum bulbocastanum is an Mi-1 gene homolog conferring broad-spectrum late blight resistance in potato: Isolation of the late blight resistance gene Rpi-blb2. The Plant Journal 44: 208–222.

van de Vossenberg BTLH, Prodhomme C, van Arkel G, van Gent-Pelzer MPE, Bergervoet M, Brankovics B, Przetakiewicz J, Visser RGF, van der Lee TAJ, Vossen J-H. 2019. The Synchytrium endobioticum AvrSen1 Triggers a Hypersensitive Response in Sen1 Potatoes While Natural Variants Evade Detection. Molecular Plant Microbe Interactions 32: 1536–1546.

van der Vossen E, Sikkema A, Hekkert B te L, Gros J, Stevens P, Muskens M, Wouters D, Pereira A, Stiekema W, Allefs S. 2003. An ancient R gene from the wild potato species Solanum bulbocastanum confers broad-spectrum resistance to Phytophthora infestans in cultivated potato and tomato. The Plant Journal 36: 867–882.

Weber E, Engler C, Gruetzner R, Werner S, Marillonnet S. 2011. A Modular Cloning System for Standardized Assembly of Multigene Constructs (J Peccoud, Ed.). PLoS ONE 6: e16765.

Werner S, Engler C, Weber E, Gruetzner R, Marillonnet S. 2012. Fast track assembly of multigene constructs using Golden Gate cloning and the MoClo system. Bioengineered 3: 38–43.

Witek K, Jupe F, Witek A. et al. (2016). Accelerated cloning of a potato late blight–resistance gene using RenSeq and SMRT sequencing. Nature Biotechnology 34, 656–660

Witek K, Lin X, Karki HS, Jupe F, Witek AI, Steuernagel B, Stam R, van Oosterhout C, Fairhead S, Cocker JM, et al. 2020. A complex resistance locus in Solanum americanum recognizes a conserved Phytophthora effector. BioRXiv 2020.05.15.095497 doi: https://doi.org/10.1101/2020.05.15.095497

Zhang C, Liu L, Wang X, Vossen J, Li G, Li T, Zheng Z, Gao J, Guo Y, Visser RGF, et al. 2014. The Ph-3 gene from Solanum pimpinellifolium encodes CC-NBS-LRR protein conferring resistance to Phytophthora infestans. Theoretical and Applied Genetics 127: 1353–1364.

Zhu S, Vossen JH, Bergervoet M, Nijenhuis M, Kodde L, Vleeshouwers VGGA, Visser RGF, Jacobsen E. 2015. An updated conventional- and a novel GM potato late blight R gene differential set for virulence monitoring of Phytophthora infestans. Euphytica 202: 219–234.

