## Supplementary data Rpi-chc1 manuscript DM, JV for "Allelic variants of the NLR protein Rpi-chc1 differentially recognise members of the *Phytophthora infestans* PexRD12/31 effector superfamily through the leucine-rich repeat domain"

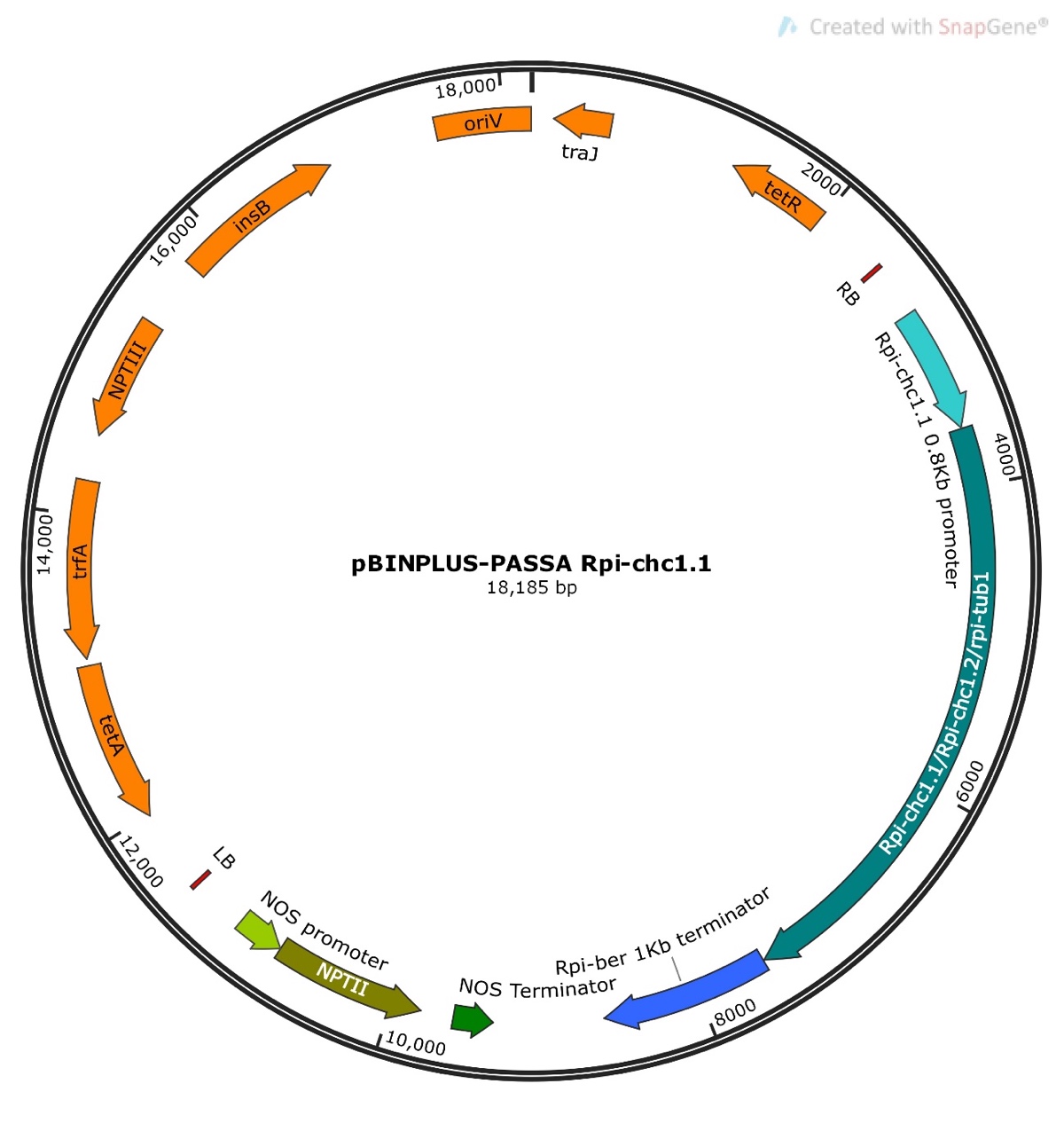

**Fig. S1**. pBINPLUS-PASSA-GG vector map. Chimeric Rpi-chc1 genes were cloned under the 0.8 Kb *Rpi-chc1.1* promoter and the 1 Kb *Rpi-ber1.1* terminator. The vector map was built using SnapGene® software (GSL Biotech).

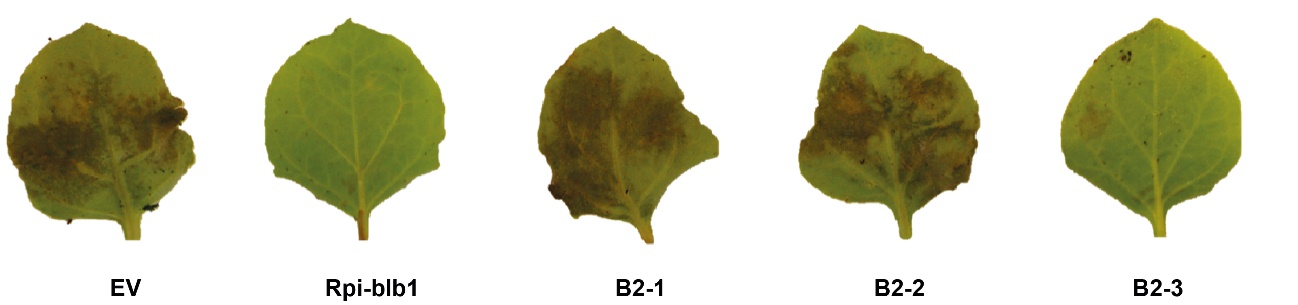

**Fig. S2**. Inoculation of P. infestans on N. benthamiana leaves agroinfiltrated with Rpi-chc1.1 candidates. The three candidates were expressed through agroinfiltration in *N. benthamiana* leaves. An empty vector (EV) and *Rpi-blb1* were used as negative and positive controls, respectively. Only candidate B2-3 was able to compromise the growth of isolate 90128.

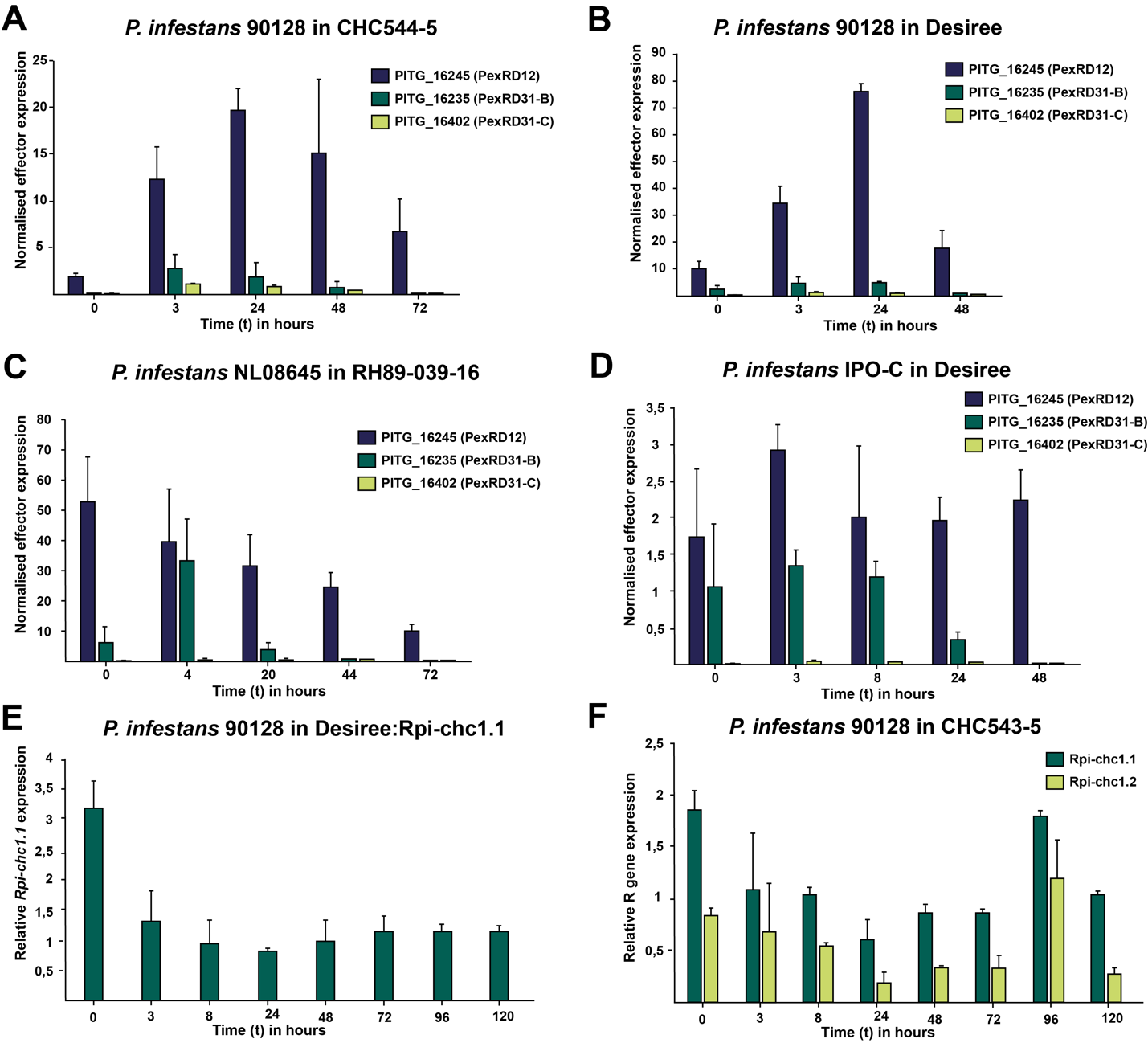

**Fig. S3.** Effector and *R* gene expression in potato leaves inoculated with *P infestans*.

(A-D) The expression of effectors PITG_16245 (PexRD12), PITG_16235 (PexRD31-B) and PITG_16402 (PexRD31-C) was evaluated in different *P. infestans* isolates after inoculation on different *Solanum* genotypes. Normalised expression data was obtained by dividing the relative effector gene expression by the *P. infestans* elongation factor 2 gene (*ef2*) expression. (E-F) *Rpi-chc1.1* and *Rpi-chc1.2* gene expression was evaluated during *P. infestans* 90128 infection. Error bars indicate the standard error of the mean (SEM).

```
1   MNYCLPSSTLQTTTKRRLTLRRLCCKQLRKKTLVQTAEEEEANTTMADPVIGATVQVLLE  60
61  KLISLTIEEVNSSRDFNKDLEMLTQNVSLIQAFIHDVETPQEKQQSVEQWLNRLERVAED 120
121 AQNVFDRFIYESLKTKVVRSPLKKVSGFFSHTAFKRKMSQKINNINKELTAINKVAKDLG 180
181 LQSLMVPSRKILPIRETDSFVVASDIVGRDLDIAEIKEKILNMREEDIVLS          231

232 TIPIVGMGGLGKTTVAKRIYNDEHMKQIFEKRIWLCLPEMSETKSFLEQILESLIERKIE 291
292 VERRDIIVKKLQDELGGKKYLLVLDDLWCVDSTSWHEFIDTLRGINTSRGNCILVTTRRK 351
352 QVASTVATDLHILGKLTEDHCWSIFKQKAFVDGRVPEELASMGNKIVKMCQGLPLAASVL 411
412 GGLLHNKEKHEWQAILDGNLLVAGEDDNGENSIKKILKLSYDYLPSPHLKKCFAYFAMFP 471
472 KDYMFEKDQLIQLWMAEGFLRPSQEIPVMEDVGHRFFQILLQNSLLQDVLLDEHNNITHC 531
532 KMHDLVHDLAGDILKSRLFDPKGDN                                    556

557  GEKLSQVRYFGCESPTDQIDK	                                   578
579  IYEPERLCTLFWRSNYTSKDM                                       598
599  LLNFKFLRVLDLSSSGIKELSAK                                     621
622  IGKLIYLRYLDLSNTEITALPNS                                     644
645  ICKLYNLQTFRVINCFSLQELPYE                                    668
669  MRNMISLRHIYYTSVDETSGHWGGWCLHNEHFQIPLNMGQ                    708
709  LTSLQTLKFFKVGLEKGRQIEELGHLKN                                736
737  LRGELTINGLQLVCDKEEAQTAYLHDKPN                               765
766  ICKLAYLWSHDESEGCEINDEHVLDG                                  791
792  LQPHPNLKTLAVVDYLGTKFPSWFSEES                                819
820     LPNLVKLKLSGSKRCKEIPS                                     839
840  LGQLKFLRHLELIGFHELECIGPAFYGVEMRNIGSNSI                      877
878  IQVFPSLKKLVLKDMRSLIEWKGDEVG                                 904
905  VRMSPGLEKLRITDCPLLKSIPNQ                                    928
929     FEILRQLKITGVDSEMPLLNLCSN                                 952
953     LTSLVKLRVYDMKELTCLPDEM                                   974
975  LRNNVSLQQIIIFNCGEFRELPQS                                    998
999  LYNLHSLRRLDIYNCTNFSSLPVPNG                                 1024
1025 DNYLTSLEFFCLHNCNGLISIPIG                                   1048
1049 MLDQCRLVFLNVSCCNNLVSFPVH                                   1072
1073 VWEMPSLSYLLISECPKLISVPKVG                                  1097
1098 LHHLTGLVRLGIGPFSEMVDFDAFQLIFNG                             1127
1128 IQQLLSLRDLEVYGRGHWDSLPYQ                                   1151
1152 LMQLSDLREITIADFGIEALPPT                                    1174
1175 LDNLTSLESLTLVRCKQLQHLNF                                    1197
1198 SDAMPKLRLLWIRDCPLLEALSDG                                   1221
1222 LGNLVSLEELYLHDCEKLEHLPSRDA                                 1247
1248 MRRLTKLWNMRIKGCPKLEESFTNYSQ                                1274
1275 WSKISHISNIELGGWRRTAISLGFSFTF                               1302

     LxxLxxLxxLxLxxCxxLxxxP
```

**Fig. S4**. Rpi-chc1.1 protein domain organization.

The N-terminal CC-domain comprises amino acids 1-231. The amino acids depicted in green shading are predicted to fold into a coiled coil structure using the “coil” algorithm with window size 14. The 21 underlined amino acids represent the MADA motif present in all Rpi-chc1 alleles. Residues in bold match the consensus MADAxVSFxVxKLxxLLxxEx. Next, 7 amino acids are underlined that match the consensus hhGRExE. The central NB-ARC domain comprises amino acids 232-557. Amino acids in red shading show similarity to the Kinase 1a, Kinase 2, kinase 3a, GLPL, RNBS-D and MHD domains, respectively. The C-terminal LRR-domain consists of 29 imperfect leucine rich repeats. The consensus sequence LxxLxxLxxLxLxxCxxLxxxP is indicated below. Conserved hydrophobic amino acids (A, V, L, and F) are marked by pink shading. LRR 3 and 4 contain the conserved VLDL motifs (underlined).

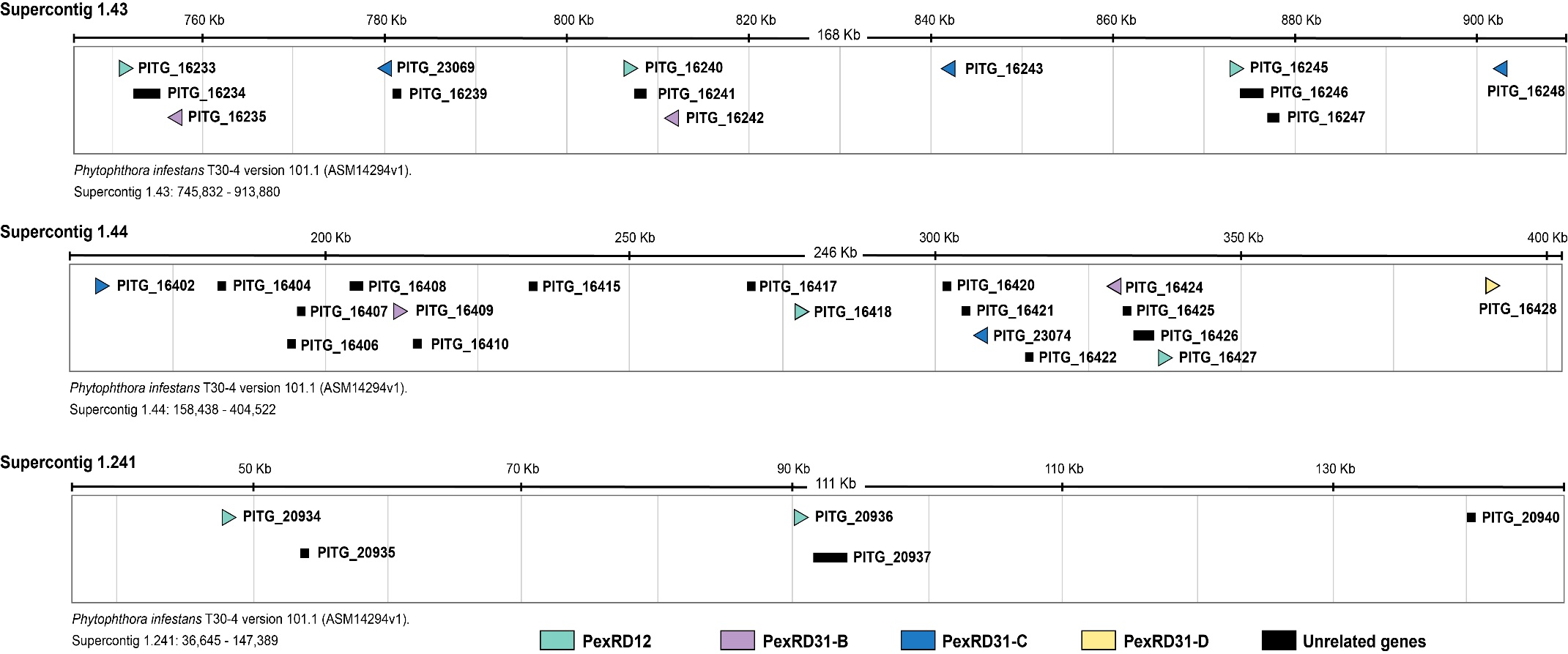

**Fig. S5**. Localization of PexRD12/31 effectors in the P. infestans T30-4 contigs. Position of the PexRD12/31 family members on genomic supercontigs of *Phytophthora infestans* T30-4 version 101.1 (ASM14294v1). PexRD12, PexRD31-B, PexRD31-C and PexRD31-D are represented in green, purple, blue and yellow, respectively. Unrelated genes have been indicated in black. Gene direction is represented by the direction of the triangles.

**Table S1.** Accession numbers of *Solanum* genotypes and *Rpi-chc1* sequences.

| **Seedling name** | **Solanum species** | **Country of Origin** | **Stock center Accession numbers** | **Rpi-chc1 alleles** | **GenBank Accession numbers** |
| --- | --- | --- | --- | --- | --- |
| 94-2031 | *S. berthaultii* | Bolivia | PI473331 | Rpi-ber1.1_94-2031 | MW390806 |
| 324-2 | *S. berthaultii* | Bolivia | CGN 18042; BGRC 18548; GLKS 0161 | Rpi-ber1.1_324-2; Rpi-ber1.2_324-2 | MW410790; MW410793 |
| 487-1 | *S. berthaultii* | Bolivia | CGN 20645; PI 498104; BGRC28012; GLKS 5433 | rpi-ber1.3_487-1 | MW410798 |
| 493-5 | *S. berthaultii* | Bolivia | CGN 17823; PI 265858; BGRC 10063; GLKS 1620 | Rpi-ber1.2_493-5 | MW410794 |
| 493-7 | *S. berthaultii* | Bolivia | CGN 17823; PI 265858; BGRC 10063; GLKS 1620 | Rpi-ber1.2_493-7;  rpi-ber1.4_493-7 | MW410795;  MW410802 |
| 493-9 | *S. berthaultii* | Bolivia | CGN 17823; PI 265858; BGRC 10063; GLKS 1620 | Rpi-ber1.2_493-9;  rpi-ber1.4_493-9 | MW410796;  MW410801 |
| 543-5 | *S. chacoense* | Bolivia | CGN 24852; PI 653755; BGRC 63055; GLKS 4994 | Rpi-chc1.1_543-5  Rpi-chc1.2_543-5 | MW383255; MW410797 |
| 544-5 | *S. chacoense* | Bolivia | CGN 18365; PI 653756; BGRC 63056; GLKS 4963 | NA | NA |
| 561-2 | *S. berthaultii* | Peru | BGRC 55178; GLKS 2798 | Rpi-ber1.1_561-2;  rpi-ber1.4_561-2 | MW410791;  MW410803 |
| 852-5 | *S. tarijense* | Bolivia | CGN 22729; BGRC 27304 | Rpi-tar1.1_852-5;  rpi-tar1.3_852-5 | MW390807;  MW410799 |
| RH89-039-16 | *S. tuberosum* | Netherlands | Wageningen U&R, Plant Breeding | rpi-tub1.3_RH89-039-16 | MW410800 |

**Table S2.** Primers used in this study.

| **Primer name** | **Primer sequence** |
| --- | --- |
| Rpi-ber Promoter Fw | CCAGTGAATTGTTAATTAAATAGG |
| Rpi-ber Promoter Rv | CTACATACCGGTATACAATCATTCAAACAGTAATAAAA |
| Rpi-ber Terminator Fw | CTACATCCTAGGGTCGCTTGCATTTTTAATTAG |
| Rpi-ber Terminator Rv | GAAACAGCTATGACCATGATTAC |
| Rpi-chc1 ORF Fw | ATGAATTATTGTCTTCCTTCG |
| Rpi-chc1 ORF Rv | TCAGAAAGTGAAAGAGAAACC |
| Chimeric LRR-2 Fw | GCTAGGTCTCGGAGAAATGTTGTTACGTAGTATCTC |
| Chimeric LRR-2 Rv | GCTAGGTCTCGTCTCTTCAACAGATAACGATTTTC |
| Chimeric LRR-8 Fw | GCTAGGTCTCGTTCACGAAACTCTCCACATTC |
| Chimeric LRR-8 Rv | GCTAGGTCTCGTGAATTGCCACAAAGCTTG |
| Chimeric LRR-14 Fw | GCTAGGTCTCGGGTGCAGTTGTAAATGTCTAATCTATA |
| Chimeric LRR-14 Rv | GCTAGGTCTCGCACCAACTTCAGTTCTCTTCCTG |
| Chimeric LRR-19 Fw | GCTAGGTCTCGCTGATCAATCCATTACAATTATGTAAG |
| Chimeric LRR-19 Rv | GCTAGGTCTCGTCAGTATACCAATTGGAATGCTAG |
| Chimeric LRR-23 Fw | GCTAGGTCTCGAGGGCAGAGAATCCCAGTG |
| Chimeric LRR-23 Rv | GCTAGGTCTCGCCCTATCAGCTTATGCAACTCTC |
| Chimeric LRR-25 Fw | GCTAGGTCTCGCATCTGAGAAGTTCAGATGTTGTAGC |
| Chimeric LRR-25 Rv | GCTAGGTCTCGGATGCCATGCCCAAATTACG |
| Chimeric LRR-29 Fw | CTACATACCGGTATGAATTATTGTCTTCCTT |
| Chimeric LRR-29 Rv | CTACATCCTAGGTCAGTAAGTGAAAGAGAAA |
| Q-PCR Rpi-chc1.1 Fw | TCAACTGCGGAGAGTTTCGT |
| Q-PCR Rpi-chc1.1 Rv | AGCTGACATTCAAAAACACTAGCC |
| Q-PCR Rpi-chc1.2 Fw | TGGTGTTGAGATTAGAAATACCG |
| Q-PCR Rpi-chc1.2 Rv | AGCTGACATGCAAATACTCTAGAGAC |
| Q-PCR PITG_16245 Fw | TTTGAGAACGCATCCGATAG |
| Q-PCR PITG_16245 Rv | TCTTTCCAGCGTGGTACTTG |
| Q-PCR PITG_16235 Fw | CATTGACGACGAAGGCTAAC |
| Q-PCR PITG_16235 Rv | GAGCGACCTGTCCTTAGAGC |
| Q-PCR PITG_16402 Fw | AATGTTGAGGACCGAGAAGG |
| Q-PCR PITG_16402 Rv | GGCCATCTAAGCCCAAGTAG |
| Q-PCR Avrsto1 Fw | CGCTTCCACATCCAAGATTCGC |
| Q-PCR Avrsto1 Rv | CGCCCGCTCTTCGTCAGAC |
| Rpi-chc1.1 Marker Fw | ACAGATAATAATTTTCAACTGC |
| Rpi-chc1.1 Marker Rv | ATTTGGGACATTCTGATATAAG |
| Rpi-chc1.2 Marker Fw | TGGTGTTGAGATTAGAAATACCG |
| Rpi-chc1.2 Marker Rv | TGTCCTTCTCCACCCACCTA |
| B2-3 sgRNA 1 Fw | AATGCTCCTGTAATTTTTAATTGA |
| B2-3 sgRNA 1 Rv | AAACTCAATTAAAAATTACAGGAG |
| B2-3 sgRNA 2 Fw | AATGTGACATTCAAAAACACTAGC |
| B2-3 sgRNA 2 Rv | AAACGCTAGTGTTTTTGAATGTCA |
| B2-3 sgRNA 3 Fw | AATGCTCACCGGGTTAGTGAGATT |
| B2-3 sgRNA 3 Rv | AAACAATCTCACTAACCCGGTGAG |

**Table S3.** *P. infestans* effectors used in this study.

| **Effectors** | **Family** | **Class** | **GenBank accession** | **Contig** | **Size (aa)** | **Reference** |
| --- | --- | --- | --- | --- | --- | --- |
| PITG_16245 | PexRD12 | A1 | XP_002897644 | supercont1.43 | 115 | Haas *et al.*, 2009 |
| PITG_16418 | PexRD12 | A1 | XP_002897459 | supercont1.44 | 115 | Haas *et al.*, 2009 |
| PITG_16427 | PexRD12 | A1 | XP_002897467 | supercont1.44 | 123 | Haas *et al.*, 2009 |
| PITG_16233 | PexRD12 | A2 | XP_002897635 | supercont1.43 | 123 | Haas *et al.*, 2009 |
| PITG_16240 | PexRD12 | A2 | XP_002897640 | supercont1.43 | 123 | Haas *et al.*, 2009 |
| PITG_20934 | PexRD12 | A2 | XP_002895188 | supercont1.241 | 123 | Haas *et al.*, 2009 |
| PITG_20936 | PexRD12 | A2 | XP_002895190 | supercont1.241 | 123 | Haas *et al.*, 2009 |
| PITG_20336 | PexRD12 | A2 | XP_002895870 | supercont1.129 | 84 | Haas *et al.*, 2009 |
| PITG_23230 | PexRD12 | A2 | XP_002909893 | supercont1.2184 | 70 | Haas *et al.*, 2009 |
| PITG_16235 | PexRD31 | B | XP_002897637 | supercont1.43 | 129 | Haas *et al.*, 2009 |
| PITG_16409 | PexRD31 | B | XP_002897455 | supercont1.44 | 129 | Haas *et al.*, 2009 |
| PITG_16242 | PexRD31 | B | XP_002897642 | supercont1.43 | 129 | Haas *et al.*, 2009 |
| PITG_16424 | PexRD31 | B | XP_002897464 | supercont1.44 | 129 | Haas *et al.*, 2009 |
| PITG_16243 | PexRD31 | C | XP_002897643 | supercont1.43 | 119 | Haas *et al.*, 2009 |
| PITG_23069 | PexRD31 | C | XP_002897638 | supercont1.43 | 119 | Haas *et al.*, 2009 |
| PITG_16402 | PexRD31 | C | XP_002897450 | supercont1.44 | 119 | Haas *et al.*, 2009 |
| PITG_23074 | PexRD31 | C | XP_002897462 | supercont1.44 | 119 | Haas *et al.*, 2009 |
| PITG_16248 | PexRD31 | C | XP_002897647 | supercont1.43 | 119 | Haas *et al.*, 2009 |
| PITG_16428 | PexRD12/31 | D | XP_002897468 | supercont1.44 | 142 | Haas *et al.*, 2009 |
| PITG_09577 | PexRD12/31 | D | XP_002903203 | supercont1.15 | 118 | Haas *et al.*, 2009 |
| PITG_14371 | Avr3a |  | XP_002898842 | supercont1.34 | 147 | Armstrong *et al.*, 2005 |

**Table S4.** CRISPR-Cas9 targeting of Rpi-chc1.1. Stable transformants were inoculated with *P. infestans* isolates IPO-C and 90128.

| **Genotype** | ***P. infestans*** | | **Phenotype** |
| --- | --- | --- | --- |
|  | **IPO-C** | **90128** |  |
| CHC543-5 | 9 | 9 | Resistant |
| CHC544-5 | 0 | 0 | Susceptible |
| CHC543-5:CRISPR-1 | 9 | 9 | Resistant |
| CHC543-5:CRISPR-2 | 9 | 9 | Resistant |
| CHC543-5:CRISPR-3 | 2 | 4 | Susceptible |
| CHC543-5:CRISPR-4 | 8 | 8 | Resistant |
| CHC543-5:CRISPR-5 | 9 | 9 | Resistant |
| CHC543-5:CRISPR-6 | 9 | 9 | Resistant |
| CHC543-5:CRISPR-7 | 9 | 9 | Resistant |
| CHC543-5:CRISPR-8 | 4 | 4 | Susceptible |
| CHC543-5:CRISPR-9 | 9 | 9 | Resistant |
| CHC543-5:CRISPR-10 | 0 | 0 | Susceptible |
| CHC543-5:CRISPR-11 | 8 | 8 | Resistant |
| CHC543-5:CRISPR-12 | 9 | 9 | Resistant |
| CHC543-5:CRISPR-13 | 0 | 0 | Susceptible |
| CHC543-5:CRISPR-14 | 9 | 9 | Resistant |
| CHC543-5:CRISPR-15 | 0 | 0 | Susceptible |
| CHC543-5:CRISPR-16 | 6 | 6 | Resistant |
| CHC543-5:CRISPR-17 | 0 | 0 | Susceptible |
| CHC543-5:CRISPR-18 | 8 | 8 | Resistant |
| CHC543-5:CRISPR-19 | 6 | 6 | Resistant |
| CHC543-5:CRISPR-20 | 0 | 0 | Susceptible |
| CHC543-5:CRISPR-21 | 2 | 2 | Susceptible |
| CHC543-5:CRISPR-22 | 4 | 4 | Susceptible |
| CHC543-5:CRISPR-23 | 6 | 6 | Resistant |
| CHC543-5:CRISPR-24 | 0 | 0 | Susceptible |
| CHC543-5:CRISPR-25 | 0 | 0 | Susceptible |
| CHC543-5:CRISPR-26 | 0 | 0 | Susceptible |
| CHC543-5:CRISPR-27 | 4 | 4 | Susceptible |
| CHC543-5:CRISPR-28 | 2 | 2 | Susceptible |
| CHC543-5:CRISPR-29 | 9 | 9 | Resistant |
| CHC543-5:CRISPR-30 | 9 | 9 | Resistant |
| CHC543-5:CRISPR-31 | 9 | 9 | Resistant |
| CHC543-5:CRISPR-32 | 4 | 4 | Susceptible |
| CHC543-5:CRISPR-33 | 9 | 9 | Resistant |
| CHC543-5:CRISPR-34 | 2 | 2 | Susceptible |
| CHC543-5:CRISPR-35 | 2 | 2 | Susceptible |
| CHC543-5:CRISPR-36 | 9 | 9 | Resistant |
| CHC543-5:CRISPR-37 | 0 | 0 | Susceptible |

* The responses in detached leaf assays were scored from 0 (complete susceptibility) to 10 (complete resistance) (average out of 6 repeats).

**Table S5**. Late blight resistance assessment of different Rpi-chc1 alleles. Desiree plants transformed with *Rpi-chc1* alleles were inoculated with the indicated *P. infestans* isolates.

| **Transformed clones*** | **Allele names** | ***Rpi-chc1* family clade** | **IPO-C** | **90128** |
| --- | --- | --- | --- | --- |
| 94-2031_L4 | *Rpi-ber1.1_94-2031* | 1 | Resistant | Resistant |
| 852-5_E28 | *Rpi-tar1.1_852-5* | 1 | Resistant | Resistant |
| 543-5_C10 | *Rpi-chc1.1* | 1 | Resistant | Resistant |
| 543-5_C2 | *Rpi-chc1.2* | 2 | Susceptible | Susceptible |
| 493-7_G12 | *Rpi-ber1.2_493-7* | 2 | Susceptible | Susceptible |
| RH89-039-16_D3 | *rpi-tub1.3_RH89-039-16* | 3 | Susceptible | Susceptible |

* the coding sequences of these alleles were cloned between the 5’ and 3’ regulatory elements of *Rpi-chc1.1* in the plant transformation vector pDEST236 using multi-site gateway cloning. The resulting plasmids were used for stable agrobacterium mediated transformation of potato variety Desiree.

**Table S6.** Segregation of markers and late blight resistance of *Rpi-chc1.1* and *Rpi-chc1.2* allelic variants.

| **Individuals from recombinant population 7650** | ***Rpi-chc1.1***  **Marker*** | ***Rpi-chc1.2***  **Marker*** | **DLA (*P. infestans* 90128)** |
| --- | --- | --- | --- |
| 4C-A07 | 1 | 0 | Resistant |
| 4C-D01 | 0 | 1 | Susceptible |
| 4C-G12 | 0 | 1 | Susceptible |
| 4D-H01 | 1 | 0 | Resistant |
| 5A-B05 | 1 | 0 | Resistant |
| 5A-B06 | 0 | 1 | Susceptible |
| 5C-E02 | 1 | 0 | Resistant |
| 5C-H04 | 0 | 1 | Susceptible |
| 5D-F02 | 1 | 0 | Resistant |
| 5D-G05 | 0 | 1 | Susceptible |
| 5E-B02 | 1 | 0 | Resistant |
| 5E-B09 | 1 | 0 | Resistant |
| 5F-C04 | 1 | 0 | Resistant |
| 5F-D07 | 1 | 0 | Resistant |
| 6B-B08 | 1 | 0 | Resistant |
| 6B-C12 | 1 | 0 | Resistant |
| 6C-F11 | 1 | 0 | Resistant |
| 6E-H03 | 0 | 1 | Susceptible |
| 7A-E05 | 0 | 1 | Susceptible |
| 7A-F11 | 0 | 1 | Susceptible |
| 7C-C03 | 0 | 1 | Susceptible |
| 7C-D03 | 0 | 1 | Susceptible |
| 7E-H04 | 1 | 0 | Resistant |
| 7E-G06 | 0 | 1 | Susceptible |
| 7E-H06 | 0 | 1 | Susceptible |
| 8A-A01 | 1 | 0 | Resistant |
| 8B-D08 | 0 | 1 | Susceptible |
| 8C-B11 | 0 | 1 | Susceptible |
| 8C-G11 | 1 | 0 | Resistant |
| 8D-F10 | 1 | 0 | Resistant |
| 8D-F12 | 0 | 1 | Susceptible |
| 8E-F06 | 0 | 1 | Susceptible |
| 8F-H02 | 0 | 1 | Susceptible |
| 8F-A04 | 0 | 1 | Susceptible |
| 8F-E03 | 0 | 1 | Susceptible |
| 8G-D03 | 1 | 0 | Resistant |
| R Parent chc543-5 | 1 | 1 | Resistant |
| S Parent chc544-5 | 0 | 0 | Susceptible |

*PCR markers for *Rpi-chc1.1* (primers MN586+MN587) and *Rpi-chc1.2* (primers 2595F+3899R)

were used to determine the presence (1) or absence (0) of each allele in the recombinant population 7650 that was derived from 1771 individuals.

**Table S7**. Functional expression of *Rpi-chc1.1* in Desiree transgenic events correlates with responsiveness to PexRD12.

| ***Rpi-chc1.1* transformation event in Desiree** | **T-DNA copy number** | **DLA**  **(90128)** | | **Field score**  **(IPO-C)** | **PexRD12-A2 responsiveness** |
| --- | --- | --- | --- | --- | --- |
| A17-03 | 2 | 8,5 | 45 | | 0 |
| A17-09 | 5 | 9,0 | 2 | | 2 |
| A17-11 | 5 | 9,0 | 6,5 | | 2 |
| A17-12 | 2 | 7,0 | 45 | | 0 |
| A17-26 | 1 | 9,0 | 50 | | 0 |
| A17-27* | 1 | 9,0 | 2 | | 2 |
| A17-29 | 4 | 9,0 | 20 | | 1 |
| A17-30 | 4 | 8,5 | 6,5 | | 2 |
| A17-31 | 3 | 7,0 | 20 | | 2 |
| A17-33 | 1 | 6,0 | 87,5 | | 0 |
| A17-34 | 2 | 4,0 | 75 | | 0 |
| A17-35 | 3 | 4,0 | 40 | | 0 |
| A17-39 | 9 | 7,0 | 2 | | 2 |
| A17-47 | 2 | 6,0 | 25 | | 0 |
| A17-48 | >10 | 9,0 | 15 | | 1 |
| A17-49 | 6 | 9,0 | 3 | | 2 |
| A17-53 | 1 | 4,0 | 85 | | 0 |
| A17-54 | 1 | 2,0 | 92,5 | | 0 |
| A17-55 | 4 | 3,0 | 45 | | 0 |
| A17-59 | 3 | 4,0 | 30 | | 0 |
| A17-79 | 2 | 6,0 | 45 | | 0 |
| A17-80 | 2 | 6,0 | 65 | | 0 |
| A17-85 | 5 | 8,0 | 35 | | 0 |
| Desiree |  | 1,0 | 92,5 | | 0 |

The T-DNA copy number was estimated by qPCR.

The responses in detached leaf assays (DLA) were scored from 0 (complete susceptibility) to 10 (complete resistance) (average out of 6 repeats).

The field score represents the percentage of infested foliage three weeks after inoculation with IPO-C (average out of 4 repeats).

PexRD12-A2 was agroinfiltrated in each plant to score the effector responsiveness (0=no response – 2=high response) (average of four repeats).

*This event corresponds to the late blight differential as described by Zhu *et al*. (2015)
